# Hexapeptides from a mammalian inhibitory hormone activate and inactivate nematode reproduction

**DOI:** 10.1101/2021.09.28.462091

**Authors:** John E. Hart, Sharad Mohan, Keith G Davies, Ben Ferneyhough, Iain J Clarke, John A Hunt, Steve D Shnyder, Christopher R Mundy, David R Howlett, Russell P Newton

## Abstract

**Background:** Biopurification has been used to disclose an evolutionarily conserved inhibitory reproductive hormone involved in tissue mass determination. A (rat) bioassay-guided physicochemical fractionation using ovine materials yielded via Edman degradation a 14-residue amino acid (aa) sequence. As a 14mer synthetic peptide (EPL001) this displayed antiproliferative and reproduction-modulating activity, while representing only a part of the native polypeptide. Even more unexpectedly, a scrambled-sequence control peptide (EPL030) did likewise.

**Methods:** Reproduction has been investigated in the nematode *Steinernema siamkayai*, using a fermentation system supplemented with different concentrations of exogenous hexapeptides. Peptide structure-activity relationships have also been studied using prostate cancer and other mammalian cells in vitro, with peptides in solution or immobilized, and via the use of mammalian assays in vivo and through molecular modelling.

**Results:** Reproduction increased (x3) in the entomopathogenic nematode *Steinernema siamkayai* after exposure to one synthetic peptide (IEPVFT), while fecundity was reduced (x0.5) after exposure to another (KLKMNG), both effects being dose-dependent. These hexamers are opposite ends of the synthetic peptide **<u>KLKMNG</u>**KN**<u>IEPVFT</u>** (EPL030). Bioactivity is unexpected as EPL030 is a control compound, based on a scrambled sequence of the test peptide MKPLTGKVKEFNNI (EPL001). EPL030 and EPL001 are both bioinformatically obscure, having no convincing matches to aa sequences in the protein databases. EPL001 has antiproliferative effects on human prostate cancer cells and rat bone marrow cells in vitro. Intracerebroventricular infusion of EPL001 in sheep was associated with elevated growth hormone in peripheral blood and reduced prolactin. The highly dissimilar EPL001 and EPL030 nonetheless have the foregoing biological effects in common in mammalian systems, while being divergently *pro-* and *anti-fecundity* respectively in the nematode *Caenorhabditis elegans*. Peptides up to a 20mer have also been shown to inhibit the proliferation of human cancer and other mammalian cells in vitro, with reproductive upregulation demonstrated previously in fish and frogs, as well as nematodes. EPL001 encodes the sheep neuroendocrine prohormone secretogranin II (sSgII), as deduced on the basis of immunoprecipitation using an anti-EPL001 antibody, with bespoke bioinformatics. Six sSgII residues are key to EPL001’s bioactivity : **<u>MKP</u>**LTGK**<u>V</u>**KE**<u>FN</u>**NI. A stereospecific bimodular tri-residue signature is described involving simultaneous accessibility for binding of the side chains of two specific trios of amino acids, MKP & VFN. An evolutionarily conserved receptor is conceptualised having dimeric binding sites, each with ligand-matching bimodular stereocentres. The bioactivity of the 14mer control peptide EPL030 and its hexapeptide progeny is due to the fortuitous assembly of subsets of the novel hormonal motif, **<u>MKPVFN</u>**, a default reproductive and tissue-building OFF signal.

**NOTE:** Please see the end of the paper for links to ten files of supplementary information and for an independent peer review.

**KEY POINTS:**

- Synthetic hexapeptides have upregulated and downregulated nematode reproduction, following similar work in other nematode species and in fish and frogs.
- Peptides up to a 20mer have also been shown to inhibit the proliferation of human cancer and other mammalian cells in vitro and influence circulating levels of ovine pituitary hormones in vivo via intracerebroventricular peptide infusion.
- The amino acid sequence of the 14mer master peptide arose from a (rat) bioassay- guided fractionation using ovine materials, whose aim was to disclose an evolutionarily conserved inhibitory reproductive hormone involved in tissue mass determination.
- The bioactive peptides mimic the non-contiguous six-residue receptor-binding active face of a polypeptide hormone, which in mammals is a derivative of the secretory vesicle prohormone secretogranin II (SgII), taking the form of a novel, structually complex 70mer dubbed ‘SgII-70’.
- The ability has been gained to reinforce or block a default reproductive and tissue- building OFF signal, using peptide mimetics.

## INTRODUCTION

During a hunt for a postulated novel hormone that is reproductively related and tissue mass reducing (Hart, 2014), ovarian follicular fluid and systemic blood plasma from sheep (*Ovis aries*) were fractionated on the basis of bioassays in vivo (organ weight reduction) (Hart, 1999) and in vitro (suppressed proliferation in bone marrow stem cells) in the rat (*Rattus norvegicus*) (Hart, 2000). Edman degradation of an active fraction of sheep plasma yielded as the cleanest N-terminal aa sequence MKPLTGKVKEFNNI (Hart, 2008), with variants on six other occasions (Supplementary Information 1, Sequencing & Purification = S1 of 10). The sequence was synthesized as a 14mer peptide with the proprietary designation EPL001 (which designation will be used hereafter interchangeably to denote both the synthetic peptide and its sequence). Resistant to conventional bioinformatic analysis (e.g. BLAST searches) and molecular biology, EPL001 is more than an incomprehensible singularity: it is an incomprehensible plurality, having been detected multiple times using Edman degradation (S1). A logo plot is shown in Fig. 1 of the seven ovine N-terminal sequences available from the physicochemical purification campaign.

**Figure 1.**
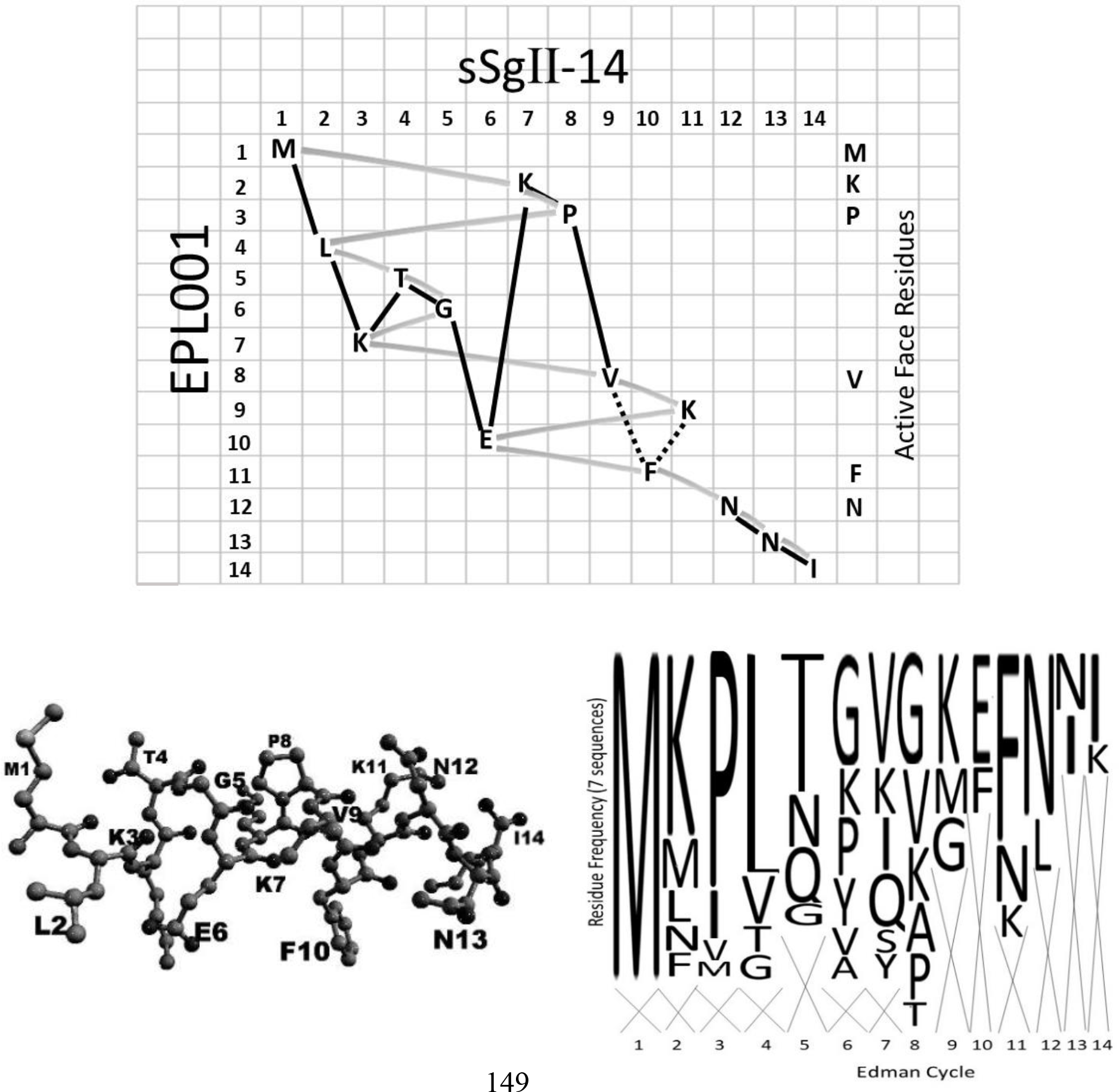
Sequence grid of sSgII-14 versus EPL001. Following the solid line yields the aa sequence of sSgII-14 across the grid, column by column. The shaded track reads out EPL001 downwards, row by row. EPL001 is interpreted as an Edman misread of sSgII-70, a hormonal proteoform of sSgII. For lysine grid placements see S2. The dotted lines are covalent bonds conjectured in the Discussion. The logo plot, lower right, is of seven available ovine N-terminal aa sequences, six of which start with M (S1). Appearing across the top of the logo plot are the proposed active face residues – in the form **<u>MKP</u>**xxx**<u>V</u>**xxx**<u>FN</u>**xx – which potentially account for the bioactivity of EPL001. Lower left is a minimized structural prediction (omitting hydrogens) for EPL143, which is sSgII-14 in contiguous form. The model takes the same general form as the grid path for sSgII-14, suggesting that the grid overall is structurally predictive.

The 14-residue N-terminal Edman sequence EPL001 was determined in the context of a polypeptide candidate of *m/z* ∼7,500 (‘Candidate 7500’) in MALDI-TOF mass spectrometry (MS) and a corresponding band in SDS polyacrylamide gel electrophoresis (Hart, 2008), implying a chain of ∼70 aa for the native molecule. In spite of comprising only 20% of Candidate 7500, EPL001 surprised by being anti-organotrophic in its own right in an assay in vivo of rat compensatory renal growth (Haylor et al, 2009) – as indeed were Candidate 7500 fractions (MS validated) of ovine ovarian follicular fluid purified by spin filtration, gel filtration and anion exchange chromatography (Hart, 2008) – and by modulating reproduction in the nematode *Caenorhabditis elegans* (Davies & Hart, 2008). Both these activities are aspects of the inhibitory hormone hypothesis (Hart, 2014). The likelihood of a project-unrelated entity having both the sought-for activities is low, especially when effects are evident in the cross- phylum manner characteristic of endocrinology. Additionally, anti-EPL001 antisera (hereafter ‘antibodies’) have provided localisation staining in mammals and in the fruit fly *Drosophila melanogaster* in immunohistochemistry (IHC) of seeming neuroendocrine significance (Hart et al, 2017). Immunostaining with anti-EPL001 antibodies was apparent in individual neurons in the ovine lateral and ventromedial hypothalamus and preoptic area, for example, with heavy staining in the palisade (neuroendocrine) region of the median eminence, with axonal beading betokening transport. The findings overall were highly reminiscent of granin protein IHC, particularly relating to secretogranin II (SgII: UniProt WEQEU8 in sheep), which is involved in the biogenesis of neuroendocrine secretory granules and is a prohormone giving rise to bioactive peptides. Further circumstantial evidence in support of the factor’s granin identity is target molecule acidity disclosed during purification (anion exchange chromatography), apparent thermostability and stoichiometric effects of semi-purified factor in assays (Bartolomucci et al, 2011). There is also a multi-way molecular weight concordance in gel electrophoresis (Hart et al, 2017). Briefly, western blots of sheep plasma deliver a single band at ∼7 kDa. This band is also seen with rat material, but a couple of other bands are seen as well, with EPL001 preabsorption in the one case tested. In the rat, the SgII gene, *Scg2* (expressed in brain, testes and adrenal: NCBI Gene ID 24765), has three exons, permitting the production of RNA splice variants. The equivalent sheep gene is mono-exonic, meaning there are no splice variants for an anti-EPL001 antibody to see.

The physicochemical purification became unproductive for unknown reasons. Candidate 7500 disappeared and thus no further aa sequence data relating to it could be obtained. An immunoprecipitation protocol was tried using an anti-EPL001 antibody to capture antigens from an aqueous extract of rat hypothalamus and fruit fly embryo material, with Orbitrap LC-MS/MS analysis after trypsinisation (Hart et al, 2017). The lead candidates to emerge from this exercise were (a) a rat protein derived by splice variation from the mammalian secretory vesicle protein secretogranin II (SgII, UniProt P10362) and (b) an SgII homologue in the fruit fly (UniProt Q9W2X8). The binding site (epitope) on EPL001 of the anti-EPL001 antibody and the epitope on any endogenous antigen were investigated using mammalian IHC as the readout (Howlett et al, 2019). This epitope mapping yielded results consistent with the mammalian endogenous antigen being SgII related and emphasized the importance of the aa motif NNI in EPL001 and in the endogenous antigen. Unconventional bioinformatics featured inclusion and exclusion criteria. Of the ovine proteome’s 26,000+ predicted proteins, the 1,100 containing NNI were included, the rest excluded. A further assumption was that a methionine residue somewhere in the ‘NNI-ome’ accounted for the initial M of EPL001, the most reliably obtained residue from seven Edman N-terminal sequence determinations obtained in two laboratories.

All M-initiated sequences in the ovine NNI-ome were eliminated that had non-EPL001 residues within 10 places of them (i.e. EPL001’s 14 residues less M and NNI: M**<u>KPLTGKVKEF</u>**NNI).

Among the few Mxxxxxxxxxx strings remaining, the stand-out candidate was one in the established prohormone, SgII (UniProt W5QEU8), the other candidates (e.g. dynein, titan) being dismissable as unlikely to deliver a secreted derivative. SgII has two main sorting domains in the form of stretches of aa directing newly synthesized SgII into intracellular secretory vesicles (Courel et al, 2008). NNI is predicted to be physically adjacent to hSgII’s first sorting domain (AlphaFold, in the absence of an experimentally determined structure in Protein Data Bank), supporting a suggestion of its relevance to sorting (Hart et al, 2017).

Meanwhile, in sSgII’s second sorting domain there is to be found 367**ML**K**TG**E**KP**V375 (with signal sequence included in residue numbering). Call this string sSgII-9. It is a lightly shuffled version of EPL001. The emphasized residues in sSgII-9 are a match for those from the front half of EPL001, in the form of three doubletons (two being exact matches to EPL001: M**<u>KP</u>**L**<u>TG</u>**KVK), interleaved by singleton matches to the second half of EPL001. The likelihood of a pair of pairs occurring by chance in two 9mer sequences is 1 in 1640 (S2, Statistics). The full complement of EPL001 residues in sSgII is represented by 367MLKTGEKPV375 and 236NNI238, leaving K and F unaccounted for to complete the anagram, defined as a rearrangement of the letters in either a word or phrase. These 14 residues constitute sSgII-14 in Fig. 1, in which sSgII-14 reads across, column by column, and EPL001 reads down, row by row. The concept here is that the EPL001 sequence is a machine artefact arising from an sSgII proteoform in the Edman machine’s reaction cartridge undergoing depolymerisation on the supporting membrane leading to aberrant sequencing. EPL001 gives the complement of aa *and* encodes their arrangement in space (see Discussion).

As a control compound there was synthesized EPL030, having the aa sequence KLKMNGKNIEPVFT. This is a scrambling of EPL001’s sequence, based on cards drawn at random by one of us (CRM). By intention a negative (i.e. inactive) control, EPL030 turned out to be contra-active in its own right in nematode studies. Investigations in *C. elegans* showed that the exogenous peptides EPL001 and EPL030 administered at 1µM per day into the aqueous medium in which this nematode can be maintained can influence the number of eggs produced (fecundity) and lifespan (Davies & Hart, 2008). The figures for fecundity were EPL001 +43%, EPL030 −64% (cumulative totals vs untreated controls, both *P*<0.001, ANOVA), with the corresponding figures for lifespan +18% and +23% (average lifespan vs untreated controls, *P*<0.005 and ns, respectively, *t*-test). EPL030’s fecundity-reducing activity has been shown to be dose-dependent and shared by another scrambled control peptide, EPL040, having a similar aa sequence (Davies & Hart, 2008). The N-terminal ‘prolinaceous sextet’ (i.e. proline-containing hexamer) of EPL001, MKPLTG (EPL016), increased the cumulative offspring of *C. elegans* by 50%, while the result for the C-terminal prolinaceous sextet of EPL030, IEPVFT (EPL036), was +79% (cumulative totals vs untreated controls, both *P*<0.02, ANOVA) (Davies et al, 2015). The reversal of the (already unexpected) *anti-fecundity* effect of EPL030 by removing its eight N-terminal aa to provide EPL036 was doubly surprising. EPL036 has also produced more offspring in molluscs, fish and frogs (Davies et al, 2015), suggesting an evolutionarily conserved system between invertebrates and vertebrates.

The hypothesis here is that peptide activity is due to an SgII-related hormonal motif; the null hypothesis that there is SgII-unrelatedness, even to the point of there being no common biological motif to explain the bioactivity of the peptides under test. Given that the prolinaceous sextets MKPLTG (EPL016) and IEPVFT (EPL036), having only a P and T in common, are both bioactive and that their parent 14mers EPL001 and EPL030 are too, there is a lot of explaining to do in terms of defining common aa residues critical for activity, for peptide optimisation. Even the C-terminus of EPL001, KVKEFNNI (EPL018), shows antiproliferative activity in vitro (Hart, 2008). The first assumption in the structure-activity analysis is that the observed effects in assays are specific, as opposed to non-specific; the second is that not all 14 residues in EPL001 are critical for activity – that there are a few key aa involved as ‘active face residues’ in binding to cellular receptors (which assumption confers analytical tractability); thirdly, given peptide diversity, that only a subset of these active face residues are required for activity; and, fourthly, that EPL001 is a form of sSgII in disguise. With a dearth of structure- activity correlates on hand, justifying the present study series, a best guess has to be hazarded as to the identity of active face residues. The assertion here is that the active face residues in EPL001 are M, K, P, V, F & N, distributed thus: **<u>MKP</u>**LTGK**<u>V</u>**KE**<u>FN</u>**NI. The conjecture is that bioactivity is evinced when these six residues or a subset are present in synthetic peptides in an appropriate conformation. The hexapeptides both have three active face residues apiece: **<u>MKP</u>**LTG and IE**<u>PVF</u>**T. The six aa have been identified on the basis of *correlated prominence* in five respects: (i) evolutionary conservation, (ii) purification prevalence, (iii) bioactivity, (iv) molecular modelling and (v) sequence gridding. A three-way doubleton match can be noted: 1**M**xxxxx**K**7 in EPL001 is present in the second sorting domain of sSgII, as 367**M**xxxxx**K**373, and in the homologue thereof in the fruit fly protein Q9W2X8 as 1048**M**xxxxx**K**1054. K373 in sSgII is followed by 374**<u>PV</u>**375, which evolutionarily conserved motif appears as 1060**<u>PV</u>**1061 in the second sorting domain of the fly protein. This doubleton is present fortuitously in the scrambled- sequence control EPL030 (KLKMNGKNIE**<u>PV</u>**FT) and in its C-terminal derivative EPL036 (IE**<u>PV</u>**FT). P and V do not appear together in EPL001, but these residues were present contiguously in the first-ever Edman reading (S1) and additionally in the sketchiest reading from the Edman campaign involving maximally purified sheep material: ultrafiltered ovine blood plasma subject also to gel filtration and anion exchange chromatography. This latter sequencing run yielded only four aa identifications interspersed with unreadable residues, with the four in register corresponding to aa in the canonical EPL001 sequence thus: MK**<u>P</u>**LTGK**<u>V</u>**KE**<u>FN</u>**NI. The minimal sequence suggests that in the source protein the four residues P, V, F & N (the ‘Beale 4’, so named for the scientist who found them) are prominent and share proximity in 3D space, as suggested by the sequence grid of Fig. 1. The prolinaceous sextets were derived empirically, as bioactive, having a position-3 proline in common. Yet preliminary molecular modelling in silico (RPN) drew the eye to EPL001’s C-terminal FNNI as a biologically relevant motif, as alluded to elsewhere (Hart, 2008), suggesting the involvement of the Beale 4’s **<u>FN</u>**. EPL001’s C-terminal KE**<u>FN</u>**NI is the epitope of the anti-EPL001 antibody (G530) and also probably the SgII-related endogenous epitope (Howlett et al, 2019), indicating surface accessibility for the antibody in both cases. The epitope residues within EPL001 are ranged in the correct order across the foot of the sequence grid. Why? Meanwhile, the proposed active face residues are ranged across the top of the grid, in the form of MKP, dipping down to VFN, with the sequence logo (Fig. 1, lower panel) portraying these six residues as among the landmarks in the physicochemical purification campaign. Layered across the middle of the sequence grid are LTG, residues deemed uninvolved in antibody or receptor binding. Besides matching up residues exogenous and endogenous, the sequence grid appears to offer a contoured functional landscape. The results of the present peptide structure-activity work arguably illuminate an evolutionarily conserved biological motif relating to a polypeptide product of SgII, of relevance to reproductive status and tissue-mass determination.

## MATERIALS AND METHODS

### Peptides

The aa sequences of 21 chemically synthetized peptides are given in Table 1. Proposed active site residues are emphasized within the hypothesis of SgII-relatedness. The peptides are in three categories: the found 14mer EPL001 and its derivatives; EPL001’s scrambled-sequence control EPL030 and its derivatives (and other 14mer control peptides); and a series of peptides modelled on EPL001 and SgII together, the EPL140s. Of these peptides, 17 are ovine-related, four human-related, the latter having hSgII’s N in place of sSgII’s V. Five 14mers have the same residues and are therefore anagrams of one another: EPL001, its scrambled-sequence controls EPL030 & EPL040 and the model peptides EPL142 & EPL143. Peptides for the nematode work were synthesized using Fmoc solid phase synthesis from l-isomer aa by Insight Biotechnology of Wembley, Middlesex, UK, and purity tested by HPLC and MS analysis as >98% pure. Peptides for the prostate cancer cell assays involving ^3^H-thymidine incorporation and the mammalian studies in vivo were synthesized at the University of Sheffield, also via solid phase peptide synthesis (Model 9050, Milligen). Peptides for all other studies were synthesized by a commercial supplier (Peptide Protein Research, Fareham, UK). Manufacture was to GLP using Fmoc solid phase synthesis from l-isomer amino acids. Purification to >98% involved RP-HPLC using water and acetonitrile as the mobile phases. Peptides were then analysed via LC-MS to determine mass and purity. Microbial contamination is precluded by the chemical nature of the processes and HPLC purification. All peptides regardless of supplier were stored in lyophilised form at −20°C prior to being taken up into aqueous stock solutions for use. Selected peptides feature in the patent literature (Hart, 2008, Hart, 2021) and EPL036 has been beta tested at third party sites under a proprietary name (www.disiaq.com).

**Table 1.** Synthetic peptides with SgII-related proposed active face residues indicated. For structure-activity purposes, interatomic distances have been measured between six atoms of the proposed active face residues using molecular models in silico. Twenty peptides provided the observed (O) distances for a chi-squared hypothesis test of SgII-relatedness, with the expected (E) interatomic distances from EPL143, a contiguous version of sSgII-14 (Fig. 1). χ^2^ = Σ (O – E)^2^/E; dof = degrees of freedom; P = probability; E-5 = x10^-5^ etc. The righthand column shows the accessibility for receptor binding of active face aa side chains, each tri-residue module viewed clockwise and counter-clockwise. Control tri-residues: **<u>MK7</u>**xx**<u>P</u>**/**<u>N</u>**x**<u>V</u>**xxx**<u>F</u>** (EPL010); **<u>K3M</u>**xxxxxx**<u>P</u>**/**<u>N5</u>**xxxxxx**<u>VF</u>** (EPL030); & **<u>K3</u>**x**<u>M</u>**xxxxx**<u>P</u>**/**<u>N4</u>**xxxxxxx**<u>VF</u>** (EPL040). NA = not applicable.

| Peptide code | Amino acid sequence and proposed active face residues: <u>M</u>•<u>KPV</u>/<u>NF</u>•<u>N</u> | Description | Interatomic distances vs sSgII-14 (EPL143) |  |  | Accessibility <u>MKP/VFN</u><br>○○/○○ |
| --- | --- | --- | --- | --- | --- | --- |
| | | | $\chi^2$ | dof | P | |
| EPL001 | <u>MKPLTGKVKEFNNI</u> | 14mer master test peptide; ovine provenance; Edman N-terminal sequence; EPL120 is extension to 20mer; EPL030 is the main scrambled EPL001 control, with EPL040 similar | 44.1 | 14 | 5.8E-5 | + +/+ - |
| EPL016 | <u>MKPLTG</u> | EPL001 Front 6 | 5.3 | 2 | 0.069 | + -/NA |
| EPL018 | <u>KVKEFNNI</u> | EPL001 Back 8 | 5.2 | 2 | 0.076 | NA/+ - |
| EPL590 | AAAAAGK<u>VKEFNNI</u> | Substituted EPL001; alanines 1-5 | 2.8 | 2 | 0.244 | NA/+ - |
| EPL545 | <u>MKPLTAAAAEFNNI</u> | Ditto; 6-9 | 31.0 | 9 | 2.9E-4 | + -/NA |
| EPL104 | <u>MKPLTGKVKEAAAA</u> | Ditto; 11-14 | 5.2 | 5 | 0.391 | + +/NA |
| EPL120 | <u>MKPLTGKVKEFNNI</u><br>KGFGVI | EPL001 Edman sequence tentatively extended 6 further reads in one ovine biopurification run, to make a 20mer | 42.2 | 14 | 1.1E-4 | + +/+ + |
| EPL010 | KGNEVM<u>KLFP</u>TINE | Scrambled control (Haylor et al, 2009) | 35.2 | 14 | 0.009 | + -/+ + |
| EPL030 | KL<u>KMNG</u>KNIE<u>PVFT</u> | Scrambled EPL001 control; MK reversal shaded | 26.3 | 14 | 0.024 | + -/+ + |
| EPL630 | KL<u>KMNG</u> | EPL030 Front 6; ditto | 4.9 | 2 | 0.088 | NA/NA |
| EPL036 | <u>IEPVFT</u> | EPL030 Back 6 (also that of EPL040) | 9.6E-3 | 2 | 0.995 | NA/NA |
| EPL040 | KL<u>KNMGNKIEPVFT</u> | Scrambled EPL001 control | 36.8 | 14 | 7.9E-4 | - -/+ + |
| EPL140 | <u>MLKTGEKPNKFNNI</u> | Variant of human model compound EPL141, with KF instead of FK | 5.9 | 9 | 0.748 | + -/NA |
| EPL141 | <u>MLKTGEKPNFKNNI</u> | Human version of EPL143, with sheep V replaced by human N9 | 0.1 | 9 | 0.999 | + -/NA |
| EPL142 | <u>MLKTGEKPVKFNNI</u> | Variant of sheep model compound EPL143, with KF instead of FK | 7.7 | 14 | 0.905 | + -/+ - |
| EPL143 | <u>MLKTGEKPVFKNNI</u> | Benchmark for chi-squared test of interatomic distances. Contiguous form of sSgII-14 (Fig. 1), an anagram of EPL001 derived from sSgII's 2<sup>nd</sup> sorting domain <sup>367</sup>MLKTGEKPV<sup>375</sup>, with <sup>178</sup>FK<sup>179</sup> and <sup>236</sup>NNI<sup>238</sup> from elsewhere in sSgII (see Discussion) | 0 | 14 | 1 | + -/+ + |
| EPL600 | <u>MKPNFN</u> | Hexapeptide version of postulated hSgII-70 active face, deduced to be mixed contiguous and non-contiguous: <sup>368</sup><u>M</u>•<sup>374</sup><u>KPN</u><sup>376</sup><u>F</u><sup>179</sup>•<sup>237</sup><u>N</u> | 9.9 | 9 | 0.357 | + -/NA |
| EPL601 | <u>MKPVFN</u> | Hexapeptide version of postulated sSgII-70 active face, deduced to be mixed contiguous and non-contiguous: <sup>367</sup><u>M</u>•<sup>373</sup><u>KPV</u><sup>375</sup><u>F</u><sup>178</sup>•<sup>236</sup><u>N</u> | 22.0 | 14 | 0.078 | + -/- + |
| EPL800 | <u>MKPNFN</u>NI | EPL600 with all of the C-terminal triplet <sup>237</sup>NNI<sup>239</sup> from hSgII-70 | 24.8 | 9 | 0.003 | + +/NA |
| EPL801 | <u>MKPVFN</u>NI | EPL601 with all of the C-terminal triplet <sup>236</sup>NNI<sup>238</sup> from sSgII-70 | 23.3 | 14 | 0.055 | + -/- - |
| EPL901 | <u>MLKTGEKPV</u> | = sSgII-9; first 9 aa of EPL143, from sSgII's second sorting domain | 2.3 | 5 | 0.809 | + -/NA |

### Entomopathogenic nematodes

The attempt was made here to replicate prior fecundity work (Davies & Hart, 2008; Davies et al, 2015) in another species of nematode, the entomopathogenic (insect slaying) *Steinernema siamkayai* Stock, Somsook & Reid 1998. Species confirmation involved standard morphological and molecular techniques (S3, Nematodes). *S. siamkayai* is a biocontrol agent of potential economic significance in global agriculture, reducing the need for pesticides. Crucial for deployment is the production of ‘infective juveniles’. These forms penetrate the larvae of insect pests, releasing a bacterium, *Xenorhabdus* spp., which ultimately kills the larvae. Stock solutions (5 mg mL^-1^) of EPL036 (IEPVFT) and EPL630 (KLKMNG) were made up and stored at −20°C. Lipid Agar plates were prepared by pouring 5 ml media in 60 mm plastic Petri dishes. The plates were inoculated with an overnight-grown bacterial culture of *Xenorhabdus* spp. (10^9^ cfu/ml) at 100 µl/plate and incubated at 28°C for 24h. *Xenorhabdus* bacteria are symbiotically associated with *S. siamkayai*. While multiplying, *Xenorhabdus* metabolises the constituents of the lipid agar and the resulting by-products are the food supplement for the nematodes that triggers the worms to grow and multiply. From the stock solutions, peptides were introduced individually into the centre of the Lipid Agar plates with *Xenorhabdus*lawn. Two concentrations of peptide were used, 4 and 8 µL of the 5 mg mL^-1^ stock peptide solutions (making 20 and 40 µg of peptide per plate in total, respectively, i.e. ∼6 and ∼12 µM). Each peptide solution was uniformly spread using an L shaped spreader and allowed to absorb into the agar plate for 10 minutes. Nematodes were harvested beforehand as follows. Third-stage infective juveniles (IJs) of *S. siamkayai* were used to infect fourth instar *Galleria* larvae.

Following larval death, the *Galleria* cadavers were placed on a White’s Trap to harvest fresh progeny within 5-6 days (Lunau et al, 1993, Pawar et al, 2013, Mohan et al, 2017). After storage in sterile water, these new-generation IJs were concentrated and inoculated on the bacterial plates on Day 1 at 500 and 1000 IJs /plate. The eight treatments (two inocula at two peptide concentrations) and two controls (two inocula without peptides) were replicated (x3) and incubated at 28°C. Observations of behaviour and morphology were made at intervals throughout the study, then on Day 5 (i.e. after 4 units of 24h) the nematodes were harvested. Two ml of distilled water was poured on each Petri dish and gently stirred to release the nematodes from the surface. The surface washings containing suspended nematodes were collected in a standard volume of 20 ml of distilled water. The number of mature adults (male and female) and the second-generation juveniles (J1, J2, J3 and pre-adults) were counted using a dissecting microscope (x10 magnification, 4 fields) in 0.05 ml subsamples (x3) and multiplied up to the standard volume of 20 ml.

### Prostate cancer cell assays

For assays involving 14mer peptides only human prostate cancer cell lines (LNCaP, PC3 and DU145) were obtained from ATCC (American Tissue Type Culture Collection, Rockville, MD, USA). Cells were cultured in DMEM (Dulbecco’s Minimal Essential Medium, Life Technologies, Grand Island, NY, USA) supplemented with 10% fetal calf serum (FCS, heat-inactivated), 100 IU/ml penicillin and 10 μg/ml streptomycin (all CSL, Parkville, Victoria, Australia) in 75 cm^2^ culture flasks (Becton Dickinson, Franklin Lakes, NJ, USA) and were maintained at 37℃ in a humidified incubator with 5%/95% CO2/O2. Incorporation of ^3^H-thymidine was used to assess DNA synthesis. Cells were seeded overnight at 0.5 x 10^5^ cells per well in 24-well tissue culture plates in DMEM culture medium (supplemented as above with 10% FCS, penicillin and streptomycin). Cell growth was synchronised by culturing cells in DMEM containing 0.1% bovine serum albumin (supplemented with penicillin and streptomycin, but without FCS) for 24h. Cells were then exposed via fresh medium (with FCS) to EPL001 or EPL030 to achieve a final concentration of 0.06 or 6 μM, or to the aqueous vehicle (controls), for 24h, before being pulsed for 48h with 1 μCi/ml ^3^H-thymidine (Amersham International, Sydney, Australia). Data are scintillation beta counts, mean +/− SD of the average values from three independent experiments performed in triplicate (n = 9), with statistical analysis by two tailed *t*-test.

Assays involving a range of peptides from 14mer down to 6mers assessed total viable PC3 or PNT2 cells using a 96-well plate MTS colorimetric assay (CellTiter 96 Aqueous One Solution Cell Proliferation Assay, Promega, Southampton, UK), with absorbance measured at 492 nm. Cells were seeded at 8,000 per well and grown for 27h. Peptides were administered at the outset to a final concentration of 30 µM (in 2 µl) to samples in triplicate. (A higher dosage was used than in the 14mer peptides study to shorten the timecourse of response.) Controls received buffer. Data are OD mean +/ − SEM, with statistical analysis by two tailed *t-test*. A time course study of the 14mers on PC3s, using the same technique, involved measurements at 24h and 48h.

Further assays on human prostate cancer cells were conducted at a contract research organisation (Epistem, Manchester, UK). LNCaP or DU145 cells seeded at 3,000 cells/well across a 96-well plate were cultured in RPMI (phenol red free), 10% FCS, 1% sodium pyruvate, Pen/strep and L-glutamine for 24h before exposure to peptides, at a single dose level in the case of LNCaPs or multiple dose levels in the case of DU145s. (For dose levels see Results.) Controls received vehicle (1% PBS). For the DU145s peptide administration was repeated after 24h. 72h after the first administration (DU145s) or only administration (LNCaP) total viable cell numbers were measured colorimetrically (MTS) using a CellTiter 96 AQueous One Solution Cell Proliferation Assay (Promega, Southampton, UK), following manufacturer’s instructions. Briefly, the media on the plates was replaced with the CellTiter reagent diluted 1:5 in DMEM. The plates were returned to the incubator for 3 hours before reading absorbance at 492 nm on a FLUORstar OPTIMA (BMG Labtech). CellTiter data are the absorbance minus background values (CellTiter plus DMEM), expressed as (OD mean +/− SEM). Statistical analysis was by ANOVA, Dunnett’s multiple comparisons test.

### Bone marrow cell assays

The method has been described previously at length (Hart et al, 2017). Bone marrow cells, consisting of various cell types including mesenchymal stem cells and fibroblasts, were cultured from adult male rats (*Rattus norvegicus*, SD strain). These cells were passaged from a growing primary stock in a T75 tissue culture flask (Sarstedt) into a 48 well tissue culture plate (IWAKI), with a cell density of 1 x 10^5^/ml. The cell culture media, henceforth referred to as ‘supplemented a-MEMS,’ comprised α-MEMS media supplemented with 10% FCS and 1% penicillin/streptomycin (all Life Technologies, Carlsbad, CA, US). On the day of peptide addition to the cells, the supplemented α-MEMS was gently removed and fresh supplemented α-MEMS added, with 10% test material by volume to achieve micromolar concentrations, untreated controls receiving just fresh supplemented α-MEMS. (For dose levels see relevant figure legends.) All samples were filter sterilised using a Millex-GV 0.22 mM syringe filter unit (Merck, Kenilworth, NJ, USA). The cells were imaged for 24 h using an incubated Zeiss Axiovert 135M microscope with a Prior Proscan II motorised stage, at 37°C with a 5% CO2 in air feed. Each well of the 48-well tissue culture plate was then imaged for 24h every 5 min using Metamorph software (MDS Analytical Technologies, Wokingham, UK), with each well having four fields of view. The viability and cell numbers were reviewed by analysing the images captured over the 24h period and cell counts were taken at selected time points using the Metamorph software.

### Mammalian studies in vivo

A sheep study involved randomly selected Corriedale ewes (*Ovis aries*) of typically 5-6 years of age maintained under natural conditions at the Monash University Sheep Facility, Werribee, Victoria, Australia. EPL001 and EPL030 were subject to intracerebroventricular (ICV) infusion separately in vivo via a permanent cannula into the third ventricle of the brains of one sheep each at 100 µg/h for 24h (method: Barker-Gibb et al, 1995, Walsh & Clarke, 1996). The zone that forms the floor of the third ventricle is that part of the hypothalamus called the median eminence. This is the anatomical interface between the brain and the anterior pituitary.

Immunostaining in the ovine median eminence, in the form of axonal beading, has been reported with an anti-EPL001 antibody (Hart, 2017). The aim here was to evaluate the impact on the ovine hypothalamic-pituitary axis of infusion from above the hypothalamus. Circulating levels of the anterior pituitary hormones LH (lutenising hormone), GH (growth hormone) and PRL (prolactin) were measured by radioimmunoassay (RIA) prior to and in the closing stages of infusion. Systemic blood samples from an indwelling jugular venous cannula were taken at 10 min intervals for 6h prior to infusion (‘Before’) and over the last 6h of infusion (‘After’). (See S9 for the checklist, ‘Arrive 2.0 Sheep’.)

For toxicity studies female Balb/c immunodeficient nude mice were used (B & K Universal, Hull, UK), aged 6-8 weeks old. They were housed in cages in isolation cabinets in an air-conditioned room with regular alternating cycles of light and darkness and received Teklad 2018 (Envigo) diet and water *ad libitum*. EPL001 and EPL030 in PBS at pH7.4 were administered to groups of 2 mice intraperitoneally (i.p.) daily on Days 0-4 at 500 mg/kg/dose and then 1000 mg/kg/dose. Following treatment, body weight was measured on a regular basis, and behaviour and general appearance monitored visually to assess for deleterious effects (e.g. dehydration, impaired mobility, hunched posture, low body temperature, ulceration and significant body weight loss), with any effects during the study recorded. If body weight loss was >15% over a 72-hour period or if animal behaviour and appearance were significantly altered, then mice were immediately sacrificed by cervical dislocation. If no deleterious effects were seen after at least 16 days of study, then the animals were sacrificed, with the dose considered non-toxic.

The MCF-7 human breast adenocarcinoma model was selected for xenograft studies. EPL001 inhibits the proliferation in vitro of MCF-7 cells stimulated to divide by insulin-like growth factor 1 (IGF-1)(Hart, 2008). An oestrogen pellet was implanted subcutaneously in the dorsal area 24 hours prior to 2-3mm^3^ fragments of MCF-7 tumour taken from donor tumours transplanted subcutaneously in the abdominal flanks of the efficacy study mice. Once tumour volumes reached approximately 32 mm^3^ (as measured by calipers, designated treatment Day 0) mice were randomised into one of three groups (n=8) as follows: EPL001 or EPL030 at 100 mg/kg/day administered i.p. on Days 0-15, plus a peptide-untreated (PBS only) control group. Tumour volume, using calipers, and animal body weight were recorded throughout the experiment and normalised to the respective tumour volume on the initial day of treatment (Day 0). Mann-Whitney U tests were conducted to determine the statistical significance of any differences in growth rate (based on tumour volume doubling time) between control and treated groups. (See S10 for the checklist, ‘Arrive 2.0 Mouse’.)

### Peptide immobilisation studies

Could the antiproliferative activity in vitro of EPL001 in solution be more predictably evoked and investigated more readily if the peptide is instead tethered to the cell culture’s substratum? Osteogenic peptides in one experiment in vitro were markedly more active in surface-immobilised form than when provided in solution (Lee et al, 2010), presumably due to a more enduring exposure of the cells. In the first study EPL001 was immobilised by either its N terminus or C terminus and human stem cells exposed accordingly. This followed a confirmation of dissolved EPL001’s antiproliferative activity in the same laboratory (JAH). In the second study, aa sections of EPL001 were substituted with alanine strings prior to tethering, to define a functional biological motif. Into the base of each well of a 24-well plate was inserted a glass coverslip (13mm diameter, borosilicate glass, SLS, Nottingham, UK), upon the upper surface of which had been immobilised an individual peptide using a bioconjugate technique.

During preparation of the coverslips, the surfaces were modified with carboxyl (−COOH) or amine (−NH2) groups separately using a published method (Curran et al., 2005). The carboxyl on the coverslips or at the end of the peptide sequence were reacted with NHS in the presence of EDC, resulting in a semi-stable NHS ester, which could then be reacted with primary amines (−NH2) on the coverslips or at the end of the peptide sequences to form amide crosslinks. For the carboxyl modified coverslips, the final peptide grafted on to the surface via its NH2 terminus was in the form Glass-CO-NH-Peptide. For the amine modified coverslips, the final peptide grafted on to the surface via its COOH terminus was in the form Glass-NH-CO-Peptide. The concentration of the peptides across the glass was approximately 0.01 mmol/cm^2^. Uncoated glass was used as the reference control. Treatments were in triplicate (i.e. n = 3). The cells used were human marrow derived mesenchymal stem cells (hMSCs, Lonza, Slough, UK, cat No PT- 2507). In the EPL001-terminus study they were seeded at 10,000 cells per well; in the alanine substitution study the density was 5,000 cells per well. The cells were maintained in DMEM with 10% FSC. EPL001 and three alanine-substituted versions of it were deployed separately: MKPLTGKVKEFNNI (EPL001), **<u>AAAAA</u>**GKVKEFNNI (EPL590), MKPLT**<u>AAAA</u>**EFNNI (EPL545) and MKPLTGKVKE**<u>AAAA</u>** (EPL104). The peptides in the alanine substitution study were N-terminally tethered to the substrate. Cell numbers were measured by CyQuant® (Invitrogen, Paisley, UK). Other determinations at 7 and 14 days involved flow cytometry in the form of fluorescence-activated cell sorting (FACS, using a FACSort, Becton Dickinson, Oxford, UK): viability (7-AAD FACS), proliferation (Ki67 FACS), cell cycle (Propidium Iodide FACS) and stem cell phenotype (antigen expression FACS: CD34, CD45, CD31, CD105, CD90 & CD73). Statistical analysis was by ANOVA using SPSS software.

### Molecular modelling

Models in silico were developed using Molecular Modelling Pro Plus, version 6.22, and ChemSite, version 5.10, produced by ChemSW (Accelrys Inc., San Diego, USA). Models of all 21 peptides in Table 1 were constructed by sequential additions of amino acid residues. Each model was adjusted in conformation to minimize energy levels: energy minimization was carried out in 1-fs time steps, to a total of 10,000 fs, with 100 equilibrium steps per iteration.

Iterations were continued until six repeat iterations yielded no change in energy gradient. In the first instance analysis of interatomic distances involved identifying 21 atoms in the amino acid side chains and the peptide backbone of the master test peptide EPL001 and designating these atoms a-u. A subset of six of these was then selected – within the hypothesis of SgII-relatedness and granted that protein interactions primarily involve aa side chains – relating to atoms in the side chains of the proposed active site residues M (= m), K (= a), P (= p), V (= q), F (= e) and N (= i). These comprise the carbon, nitrogen or sulphur atoms in the relevant amino acid side chains most remote from the peptide bond. As such, these atoms would be free to show the most intramolecular movement thence variation in subsequent interatomic distances after energy minimization. The distances of each of these atoms was measured from the others, yielding 15 measurements in all for EPL001, in angstrom units (Å). These distances were then measured in each of the other peptides of Table 1, as appropriate, given differing aa compositions. Distances between pairs of atoms were computed automatically after atoms were selected manually on-screen. Each measurement was repeated twice more after closing the model and reloading to verify the initial measurement. Twenty peptides provided the observed (O) distances for a chi-squared hypothesis test of SgII-relatedness, with the expected (E) interatomic distances from EPL143, a contiguous version of sSgII-14 (Fig. 1). The χ^2^ formula is Σ (O – E)^2^/E. This was solved online (https://goodcalculators.com/chi-square-calculator/), with double-checking online (https://www.danielsoper.com/statcalc/calculator.aspx?id=11) and manually. Complementing the dimensional comparison is an analysis of tri-residue topology: *tri-residue*, because stereospecificity requires a minimum of three points of attachment (Easson & Stedman, 1933; Ogston, 1948), a number conferring analytical tractability in the present context; and *topology*, used here to describe structural arrangements in space in relation to side chain accessibility for receptor binding. In brief, molecular models in silico were rotated to provide 3D views of sets of three aa residues (notably MKP & VFN), read in the order of the primary peptide sequence and viewed in clockwise and counter-clockwise directions. Each of the tri-residue views was then assessed visually as to whether, for all three specified aa residues, there was collective availability for receptor binding or if access was blocked by other residues. In full, energy-minimized complete ball and stick models were subject to hydrogen atom deletion (for ease of visualization), with retention of overall 3D peptide configuration.

Tri-residues were read in the order in which they appeared in the peptide primary structure and viewed either clockwise or counter-clockwise. Models were slowly rotated so that two of the aa side chains in a tri-residue module were visible: the model was then further rotated to determine whether all of the atoms of the third tri-residue aa side chain could be visualised, or only partially obscured by its own atoms or those of the other two tri-residue side chains, while retaining visibility of the first two tri-residue side chains. If all three side chains could be so visualised, the module was designated accessible for receptor binding. In the event of obstruction by a non-tri-residue aa of a tri-residue aa, the module was deemed inaccessible.

Each assessment of accessibility was carried out three times (replicated twice, with model identities concealed), commencing from the original minimized complete ball and stick model. Other parameters examined were point charges (Del Re: Huckel), electrostatic surface maps, hydrophilic surface areas, solubility, partition coefficients, dihedral angles of side chains around peptide bonds, triatomic bond angles relating to the six active face residues and lipophilicity, as well as molecular volumes, surface areas and overall dimensions.

### Ethics

All experimental procedures were conducted in compliance with applicable laws, regulations and professional standards of probity and good faith, with appropriate ethical oversight. Data integrity has been maintained throughout these exploratory studies, without outlier exclusions and with appropriate recording and archiving. Positive outcome and null studies are reported here, without selective reporting, in the interests of transparency and the provision of a true account via open access with open data. In the matter of sample identity, proprietary codes were favoured over explanatory designations to foster blind experimentation, with unbiased analysis and impartial supervision, all of which were aided in any case by project inscrutability, which countered confirmation bias relating to either the hypothesis of SgII-relatedness or the null hypothesis of SgII-unrelatedness. The G530 goat antiserum (S4) was raised in compliance with the Australian Prevention of Cruelty to Animals Act 1986, with procedures approved by the relevant Animal Ethics Committee. The ER87 rabbit antiserum (S4) was raised at Eurogentec, Belgium. The M4 mouse monoclonal antibody (S5) was raised in the UK in compliance with relevant regulations, including the Animals (Scientific Procedures) Act 1986. Prostate cancer cell assays involving ^3^H-thymidine incorporation were conducted at Monash University, Australia, with all other assays in vitro conducted in the UK. All cell assays were performed in compliance with relevant regulations. The bone marrow cell assay work was conducted under UK Home Office Project Licence PPL 30/2280 and Personal Licence 30/353 and while specific ethical approval was not required the research was performed in accordance with the guidelines of the Ethical Review Board of the University of Reading, UK, and within its purview.

Husbandry and termination in the sheep study were carried out according to the guidelines established by the Australian Prevention of Cruelty to Animals Act 1986 and procedures were approved by the relevant Animal Ethics Committee of Monash University, Australia (approval number MARP/2012-012). The maximum tolerated dose evaluations and xenograft studies were conducted in accordance with ethical standards approved by the Animal Welfare Ethics Review Board at the University of Bradford, and in accordance with the UK National Cancer Research Institute Guidelines for the Welfare of Animals. Throughout the studies, all mice were housed in air-conditioned rooms in facilities approved by the United Kingdom Home Office to meet all current regulations and standards. All procedures were carried out under a Project Licence (PPL 40/2585) issued by the UK Home Office according to government legislation. In all studies in vivo, as few animals were used as possible, in accordance with ‘Reduction’ in the Three Rs of Replacement, Reduction, Refinement.

## RESULTS

### Entomopathogenic nematodes

There was a lack of overt toxicity across all groups of nematodes as evinced behaviourally and in terms of gross morphology. Population growth over 4 days can be seen in Fig. 2; among peptide-untreated controls in terms of both adults and IJs together it was x8.2 (i.e. 4085 total individuals for the 500-worm inoculum) & x7.2 (i.e. 7175 total individuals for the 1000-worm inoculum). For the higher dose of peptide (8 µL) the comparable figures were: EPL036 peptide IEPVFT x19.3 (i.e. 9654 total individuals for the 500-worm inoculum) & x19.2 (i.e. 19171 total individuals for the 1000-worm inoculum); and EPL630 peptide KLKMNG x4.1 (i.e. 2052 total individuals for the 500-worm inoculum) & x3.2 (i.e. 3187 total individuals for the 1000-worm inoculum). Total populations in the high dose groups at termination were over twice control levels for EPL036 and at about half control levels for EPL630.

**Figure 2.**
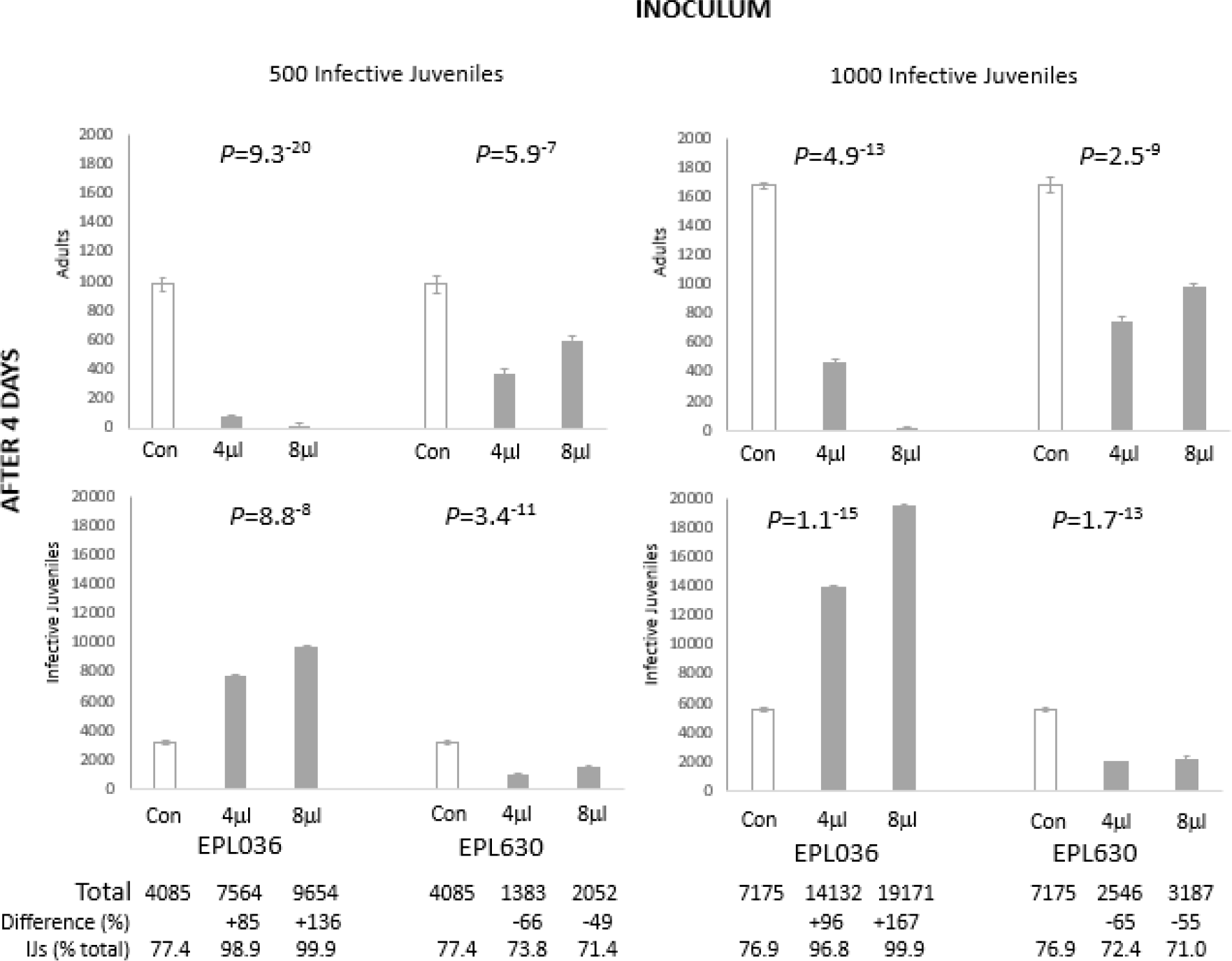
Developmental impact on *Steinernema siamkayai* of hexapeptides after 4 days. INOCULUM = founding population of infective juveniles (IJs). ‘Infective juveniles’ on the vertical axis are mostly second-generation IJs. Peptides applied at the start of the study were EPL036 = IEPVFT, EPL063 = KLKMNG. Total = Adults + Infective juveniles. Data are mean +/- SEM (average of 3 replicates). Statistical analysis: ANOVA. Control bars are doubly represented for ease of comparison. Note different vertical axes.

EPL036 induced significant increases in the rate of nematode growth and development at both inoculum rates of 500 and 1000 IJ and at both peptide concentrations, 4 and 8µg mL^-1^, dose dependently. At 4 days the majority of the first-generation EPL036-exposed IJs had transformed into mature adults which subsequently mated and went on to produce second-generation IJs (J1, J2, J3 and pre-adults), with adults barely represented, implying a die-off among these forms after an accelerated life cycle. In contrast, EPL630 suppressed to a significant degree the development of second-generation IJs, with adults at 26-29% of the population at termination, compared to controls at ∼23%. IJs in the higher dose groups at both inoculum levels were more than three times as numerous for EPL036 than controls at termination and less than half as numerous than controls for EPL630.

Observationally, infective juveniles (∼0.8 mm in length) when released on EPL036 grew and matured at a faster rate than worms in the other groups and within 24h they had transformed into pre-adults (males and females). In another 12h (i.e. 36h after incubation) they had developed into fully grown males (∼1.5 mm) and females (∼5.0 mm) and mating had commenced. In comparison, infective juveniles exposed to EPL630 or exposed to no peptide (i.e. untreated controls) transformed into pre-adults in about 36h and developed into males and females in the next 12h (i.e. 48h after incubation). With differences in development and maturation rates came differences in size. EPL036 females were bigger than those in the other two groups. They were visibly bloated with eggs compared with untreated controls, whereas egg production in EPL630-exposed females was markedly below that of controls (Fig. 3). Apart from dual inocula and dual dose levels, a duplication is provided by a pilot study, which gave similar outcomes (S3).

**Figure 3.**
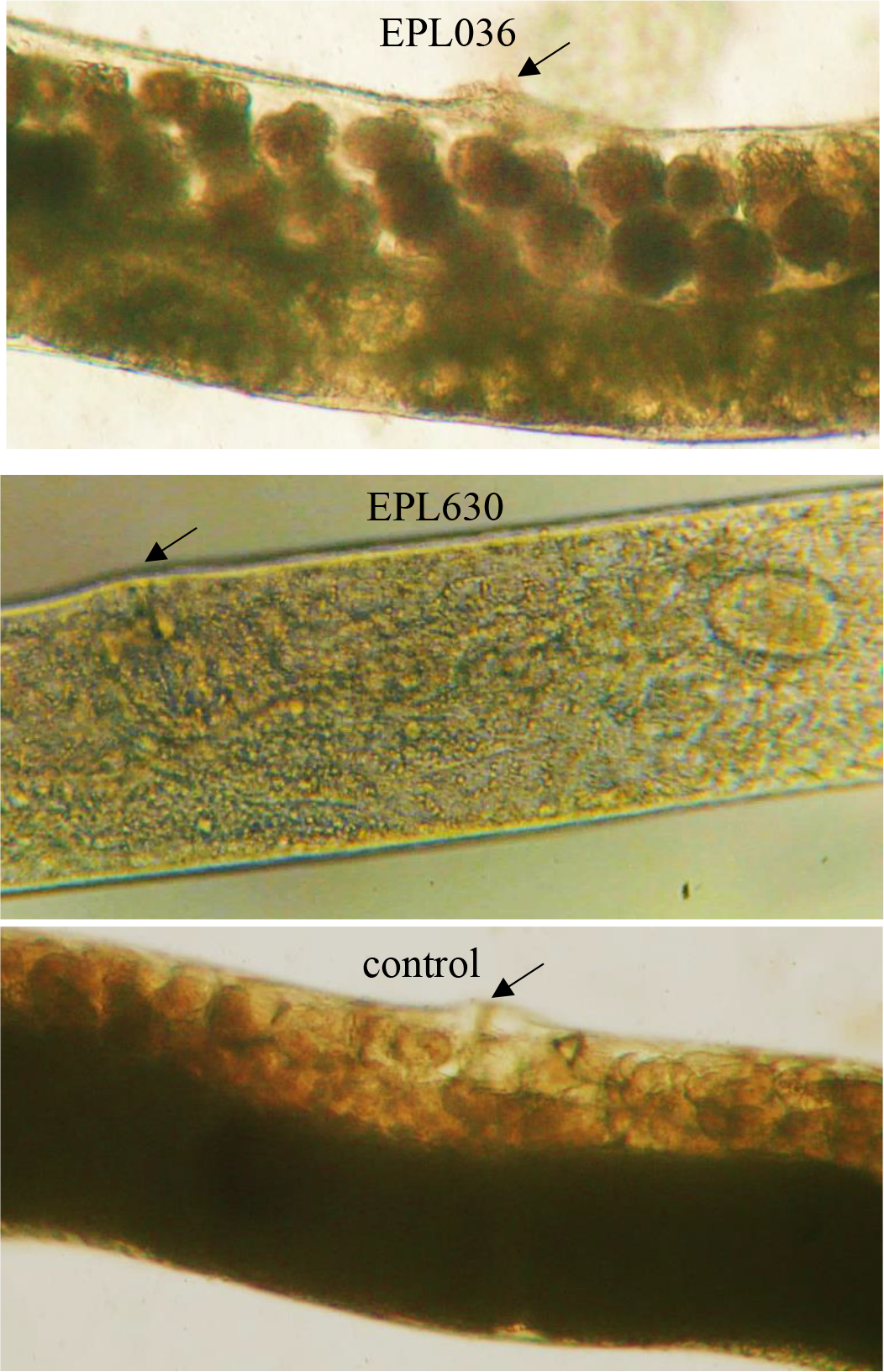
Nematode egg production. Representative mid-body photomicrographic images of adult female nematodes (*Steinernema siamkayai*) after 48h exposure to hexamer peptides EPL036 and EPL630 or unexposed to peptide (‘control’). Images obtained using a dissecting microscope at x10 magnification. Eggs appear as dark roundels in the upper image (EPL036). Fewer eggs are visible in the untreated control and fewest of all in the EPL630-exposed worm. Arrows indicate vulvas.

### Prostate cancer cell assays

Both the 14mer peptide EPL001 and its scrambled-sequence control, EPL030, were antiproliferative (Fig. 4). A dose-response relationship was more clearly evinced by EPL030 than EPL001. In the assays involving different sized peptides the results were similar for PC3 and PNT2 cells, with the smaller items associated with lower ODs than the 14mers (Fig. 5). The differences in percentage terms between test ODs and control means were: EPL001, −0.7/+3.1 (PC3/PNT2); EPL141, −22.6/−27.9; EPL143, −10.2/−20.6; EPL901, −21.2/−28.9; EPL600, −27.3/−39.7; and EPL601, −28.0/−33.2. Among the 14mers the same order of inhibitory power was seen in the PC3 time course study at 24h: EPL001<EPL143<<EPL141, with the last-mentioned mean 56% of the EPL001 mean (*P*<0.01). The ODs of all three had rebounded by 48h and were indistinguishable from one another. EPL030 on the other hand showed an enduring inhibitory effect. At 24h its OD (1.57 +/−0.03) was below that of EPL001 (1.94+/−0.09) by 18.9% (*P*<0.02); at 48h the figures were 1.15+/−0.17 and 2.36+/−0.50, respectively, with EPL030 below that of EPL001 by 51.3% (*P=*0.08, ns). For LNCaP cells over 72h, reductions in test mean ODs versus the vehicle control mean were: EPL001, -17.0%, EPL590 (= alanine substituted EPL001, **<u>AAAAA</u>**GKVKEFNNI), −21.4%, EPL600, −35.1% and EPL601, −16.0% (all *P*<0.05, with EPL600 at *P*<0.01). For DU145 cells, with peptides at 0.015-150 µM and lower cell seeding density, test mean ODs were mostly below the control mean for EPL001, EPL141, EPL143, EPL800 and EPL801, but statistical significance was only attained for EPL001 at the highest concentration (−16.5%, *P*<0.02) and EPL143 at 0.15 & 150 µM (−4.3%, *P*<0.05 and −18.7%, *P*<0.002). The 6mers evinced a degree of dose dependency, returning similar profiles. **EPL600**: 0.015 µM, 1.41 +/−0.08, +8.6%, ns; 0.15 µM, 1.29 +/−0.02, −0.1%, ns; 1.5 µM, 1.20+/−0.04, −8.4%, *P*<0.02; 15 µM, 1.24+/−0.03, −7.1%, ns; 150 µM, 1.20 +/−0.03, −16.9%, *P*<0.02. **EPL601**: 0.015 µM, 1.39 +/−0.07, +6.9%, ns; 0.15 µM, 1.24 +/−0.03, −4.3%, ns; 1.5 µM, 1.18+/−0.02, −10.4%, *P*<0.02; 15 µM, 1.18+/−0.04, −11.4%, *P*<0.03; & 150 µM, 1.24 +/−0.10, −14.3%, ns. Note that extending the 6mers to 8mers, by adding a C-terminal NI, effectively abolished activity.

**Figure 4.**
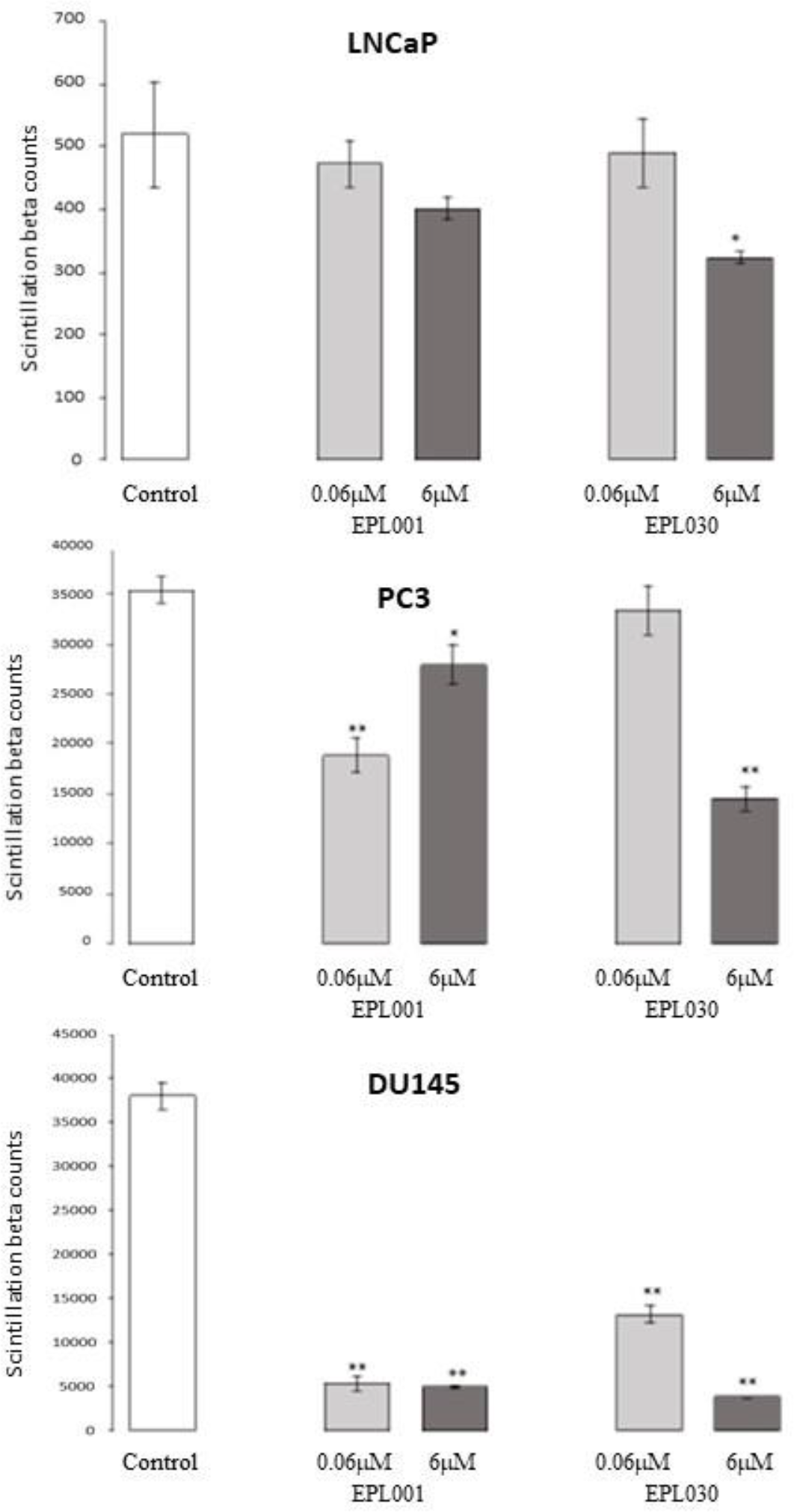
EPL001 and EPL030 effects on human prostate cancer cell growth in vitro. Cell proliferation was determined by ^3^H-thymidine incorporation. Cells were treated with the 14mer EPL001 or its scrambled-sequence control EPL030 at 0.06 or 6 μM (or given solvent control) for 24h before being pulsed for 48h with 1 μCi/ml ^3^H-thymidine. Data are scintillation beta counts, mean +/-SD of the average values from three independent experiments performed in triplicate (n = 9), with statistical analysis by two tailed *t*-test. * p<0.05; **p<0.0001.

**Figure 5.**
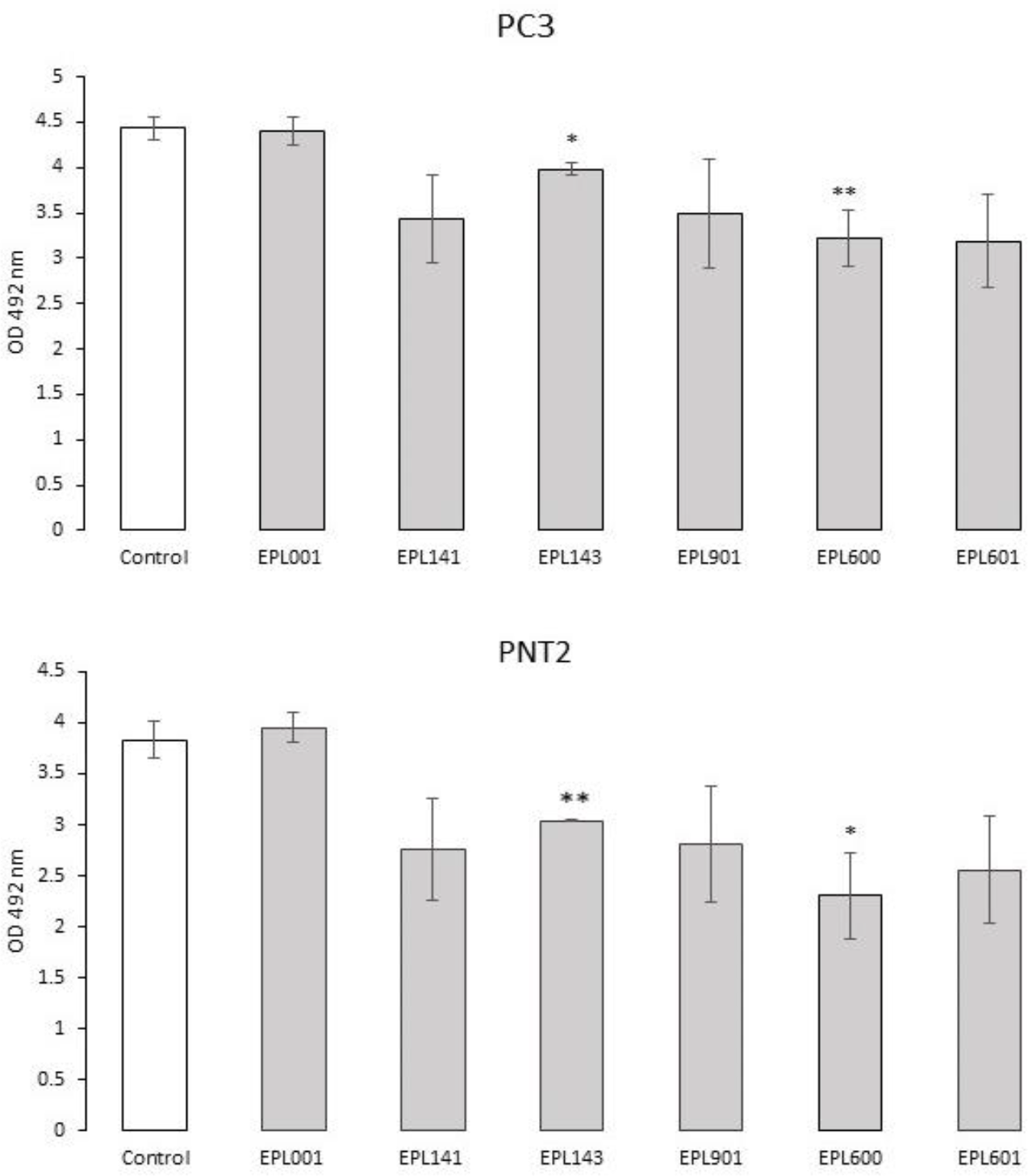
Effects of a range of peptides on human prostate cancer cell growth in vitro. Assays involved a range of peptides from 14mers down to 6mers (Table 1), with viable cells assessed using a 96-well plate MTS colorimetric assay. Absorbance measured at 492 nm. Cells were seeded at 8,000 per well and grown for 27h. Peptides were administered at the outset to a final concentration of 30 µM (in 2 µl) to samples in triplicate. Controls received buffer. Data are OD mean +/- SEM, with statistical analysis by two tailed *t-test*. * p<0.05; **p<0.025.

### Bone marrow cell assays

Whereas EPL001 was antiproliferative in this system, EPL030 was additionally pro-apoptotic (Fig. 6) as, to a more remarkable extent still, was EPL120, the 20mer extension of EPL001. The relative potencies were reversed when proline-containing hexamers from the 14mers were tried: MKPLTG (EPL016) from **<u>MKPLTG</u>**KVKEFNNI (EPL001) was a more potent inhibitor than IEPVFT (EPL036) from KLKMNGKN**<u>IEPVFT</u>** (EPL030) (Fig. 7).

**Figure 6.**
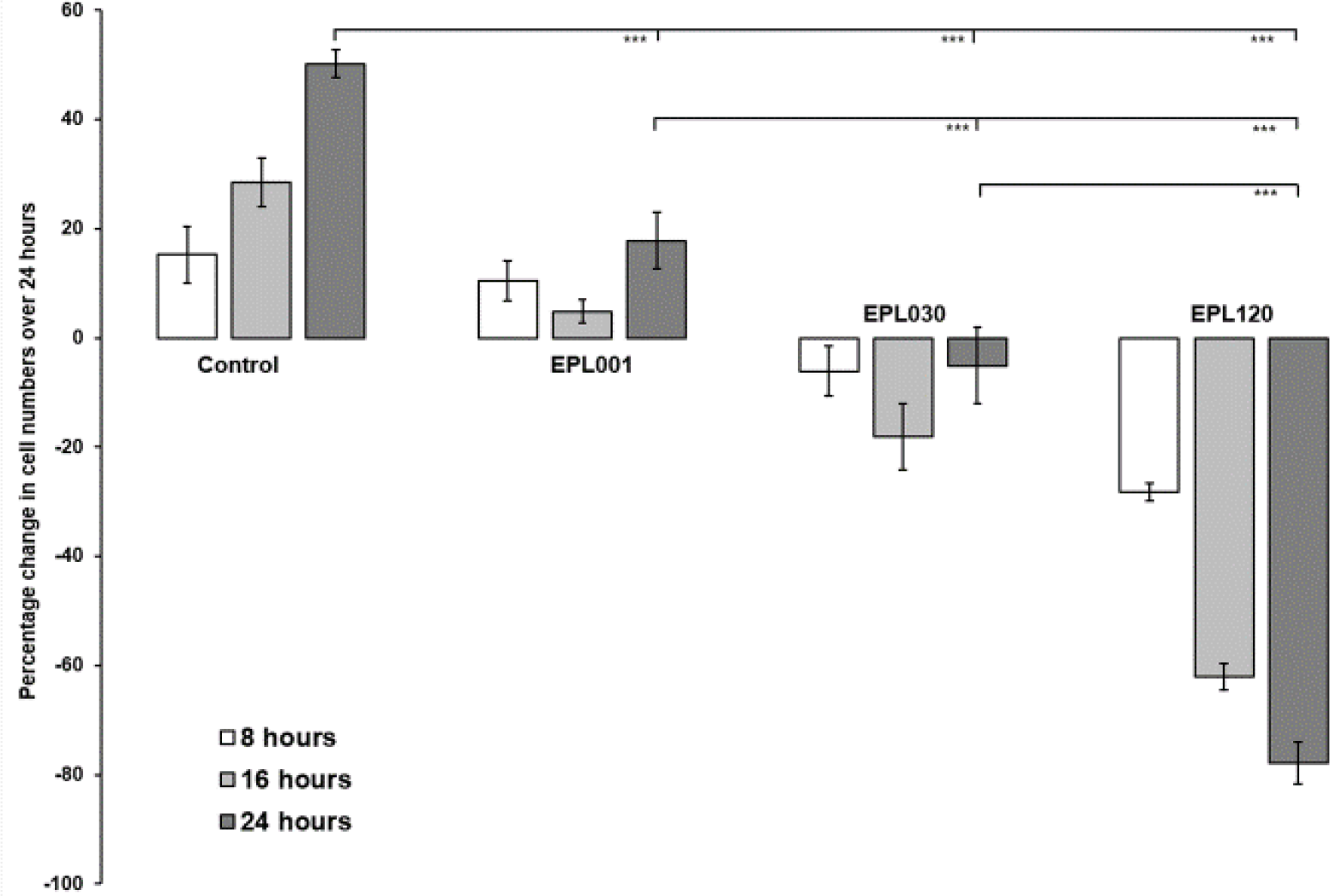
Bone marrow cells in culture exposed to long peptides. Cells were counted over 24h after exposure to the 14mer peptide EPL001 or its scrambled-sequence control EPL030 at 6µM final concentration or to the 20mer EPL120 at 5µM final concentration, compared to peptide-untreated controls.

**Figure 7.**
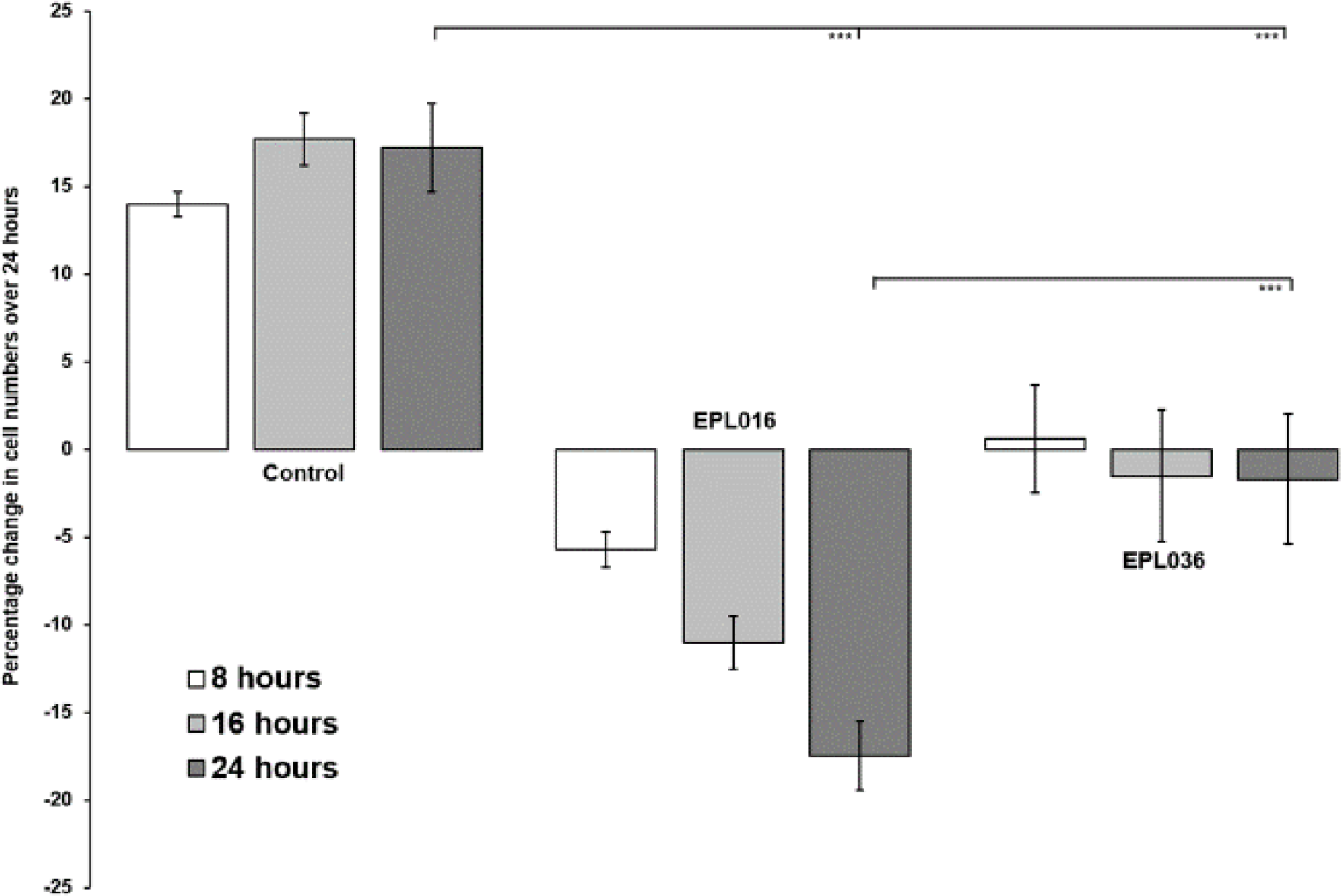
Bone marrow cells in culture exposed to hexapeptides. Cells were counted over 24h after exposure to either EPL016 (= MKPLTG, the ‘Front 6’ of the 14mer EPL001) or EPL036 (= IEPVFT, the ‘Back 6’ of EPL030, the scrambled-sequence control of EPL001) at 1µM final concentration, compared to peptide-untreated controls.

### Mammalian studies in vivo

In the ovine ICV study the RIA results in ng/ml were as follows for EPL001, Before vs After, means +/− SEM, with statistical evaluation (unpaired *t* test): **LH** 5.69 +/− 0.83 vs 6.48 +/− 0.90 (*P* = 0.521, ns), **GH** 2.72 +/− 0.60 vs 3.41 +/− 0.63 (*P* = 0.194, ns), **PRL** 132.97 +/− 16.22 vs 95.14 +/− 21.62 (*P* = 0.166, ns). The corresponding results for EPL030 were as follows: **LH** 12.00 +/− 2.89 vs 12.89 +/− 3.78 (*P* = 0.852, ns), GH 5.44 +/− 0.80 vs 7.82 +/− 0.80 (*P* = 0.039, s), **PRL** 140.54 +/− 16.22 vs 107.03 +/− 18.92 (*P* = 0.183, ns). The percentage differences as between After and Before for EPL001 were these, with the results for EPL030 in parentheses: **LH** +13.4% (+7.4%), **GH** +25.4% (+43.8%), **PRL** −28.5% (−23.8%). A vertical scattergram of these outcomes would show the same pattern for each of the peptides, with the two results for each hormone adjacent: **GH>LH>PRL**. There are six ways of arranging three items. Assuming each arrangement is equally likely the probability of selecting **GH>LH >PRL** for EPL001 is 1/6. Similarly, the probability of selecting the same order for EPL030 is also 1/6. The probability of selecting **GH>LH>PRL** for both peptides by chance is therefore 1/6 x 1/6 = 1/36 = 0.028 or 2.8%. Given that *P* = 0.028 the observed ranking arrangement **GH>LH>PRL** is unlikely to have occurred by chance. The consistency of these results from a pair of anagrammatical 14mer peptides argues against non-specific effects, though that was the initial assessment in this pilot study, halting further experiments in vivo. (See S4, Hypothalamus & Pituitary, which includes data from pituitary studies in vitro.) In the evaluation of toxicity over 14 days in mice, both EPL001 and EPL030 were well- tolerated at relatively high doses (See S5, Murine Studies). No discernible toxicity was seen with either EPL001 or EPL030 at 500mg/kg/day ip Days 0 to 4, with a non-significant maximum weight loss (compared to the initial starting weight on Day 0) of 5.0% and 4.0% respectively on Day 2. A higher dose of 1000mg/kg/day ip on Days 0 to 4 was then tried for both peptides. Again, no discernible signs of toxicity were seen, however in this case weight loss was sustained for longer with EPL030 (maximum weight loss of 7.0% on Day 3) than EPL001 (maximum weight loss of 3.5% on Day 1). The foregoing reinforces a positive safety perception in regard to the peptides under investigation (Davies et al, 2015).

In the MCF-7 tumour xenograft study, no statistically significant tumour growth delay was seen with either EPL001 or EPL030 compared to controls, although there was evidence of a slight reduction in the rate of tumour growth as the experiment progressed, more obviously with EPL030 (−20.2% at termination compared with PBS-only controls, ns)(S5 Fig. 3). It may be that increasing the administered dose to that used in the toxicity studies might have increased this effect. (A precursor study is also reported in S5, lacking peptide-untreated controls. This showed no significant differences in xenograft tumour growth between nude mice receiving either EPL001 or EPL030.)

### Peptide immobilisation studies

EPL001 was mildly antiproliferative whether tethered to coverslips N-terminally (EPL001-N) or C-terminally (EPL001-C), with EPL001-C significantly more inhibitory at 7 days. With an initial seeding of 10,000 cells per well, the results for <u>Day 7</u> were: **Glass** (i.e. controls grown on bare glass surfaces, mean cell number +/− SEM) = 46,242 +/−919; **EPL001-N** = 40,123 +/−484, −13% vs Glass; & **EPL001-C** = 32,799 +/−435, −29% vs Glass, −18% vs EPL001-N. The <u>Day 14</u> data were: **Glass** = 50,596 +/−677; **EPL001-N** = 41,314 +/−314, −18% vs Glass; & **EPL001-C** = 39,540 +/−653, −26% vs Glass, −4% vs EPL001-N. The test data points were all significantly below controls, *P*<0.05, with the EPL001-C means lower than those of EPL001-N, albeit significantly so only at Day 7 (−18%, *P*<0.05). Stem cell markers analysed by FACS were unchanged, indicating maintenance of the stem cell phenotype, and the viability measure 7-AAD was likewise unchanged. Ki67, uniform for the three groups at 7 days, was by Day 14 below the control mean for EPL001-N by 31% and below it for EPL001-C by 18%, in line with enduringly reduced cell proliferation (S6, Peptide Immobilisation & Gene Studies: Figs. 1-3 therein). The three EPL001-related peptides (N-terminally tethered) having alanines in positions 1-5 (EPL590), 6-9 (EPL545) and 11-14 (EPL104), together with EPL001-N itself, were associated with 16-29% lower mean cell numbers after 7 days than controls and 23-37% lower mean cell numbers after 14 days. With an initial seeding of 5,000 cells per well, the results for <u>Day 7</u> were: **Glass** = 14,097 +/−1,576; **EPL001-N** = 10,785 +/−2,361, −23% vs Glass; **EPL104** = 10,010 +/−4,427, −29%; **EPL545** = 10,007 +/−3,281, −29%; & **EPL590** = 11,805 +/−4,028, 16%. The <u>Day 14</u> data were: **Glass** = 26,392 +/−6,805; **EPL001-N** = 20,003 +/−5,118, −24% vs Glass; **EPL104** = 16,736 +/−1,302, −37%; **EPL545** = 18,055 +/−1,771, −32%; & **EPL590** = 20,347 +/−2,264, −23% (S6: Fig. 5b). There were no significant differences between EPL001-N and its alanine-substituted analogues, although the mean for EPL104 was below that of EPL001-N by 16% at 14 days. Compared with controls, only the data relating to EPL104 after 14 days reached statistical significance (−37%, *P*<0.05), with a higher proportion of cells for EPL104 in G0/G1 phase after 7 days (FACS analysis with PI staining, *P*<0.05) and a lower proportion in S and G2/M phases at both 7 & 14 days (also FACS analysis, *P*<0.05, corroborated by Ki67 staining, which is present during G1/S/G2/M phases only, detected in regard to 26.95% of control cells at termination versus 12.69% of EPL104- exposed cells; 18.92% of EPL001-N cells). Cells in all groups maintained their original stem cell phenotype (according to FACS analysis of CDs 29, 31, 34, 44, 45, 73, 90 & 105) and there were no effects on cell viability (7-AAD) or morphology (microscopic inspection). The alanine substitution results have been presented in fuller form via a conference poster (Pu et al, 2012, for which item see S6). Note that statistical significance for lowered cell numbers has been achieved separately in regard to the proliferation of LNCaP prostate cancer cells in vitro for EPL590 (**<u>AAAAA</u>**GKVKEFNNI); see above.

### Molecular Modelling

Interatomic distances have been determined between the proposed SgII-related active face residues M, K, P, V, F & N, along with spatial arrangements in terms of how accessible these residues are for receptor binding, via tri-residue topology. Although representing a fraction of the data that could have been generated by considering all residues and every atom (S2), the interatomic distance assessments have proved particularly productive. Apart from tri-residue topology, the other analyses (see Materials and Methods) showed no clear correlations with the relative bioactivity of the peptides. The results of a chi-squared test of SgII-relatedness are shown in Table 1, based on the analysis of 15 interatomic distances between 6 atoms (Fig. 8) within 21 peptides (S7, Interatomic Distances). The assumption here is that the geometric (steric) arrangement of atoms between the proposed active face residues is important for bioactivity. With chi-squared tests the null hypothesis is usually that the observed (O) data are the same as the expected (E) data, such that χ^2^, i.e. Σ (O – E)^2^/E, is 0. This is not the present case because the expected values are the atom-to-atom distances within an SgII-related peptide, EPL143. So χ^2^ values approaching 0, with a probability (*P*) approaching 1, uphold the hypothesis of SgII relatedness. Large χ^2^ values, with values of *P*<0.05, support to a significant extent the null hypothesis of SgII unrelatedness, in this reverse chi squared test. The benchmark for the comparison, then, is EPL143, a contiguous stand-in for the non-contiguous sSgII-14 of Fig. 1: MLKTGEKPVFKNNI. Compared with itself, EPL143 delivers a χ^2^ value of 0 (i.e. there is no difference between observed and expected), with 14 degrees of freedom (dof = n – 1 for the 15 interatomic distance measurements) and a probability, *P*, of molecular similarity of 1.

**Figure 8.**
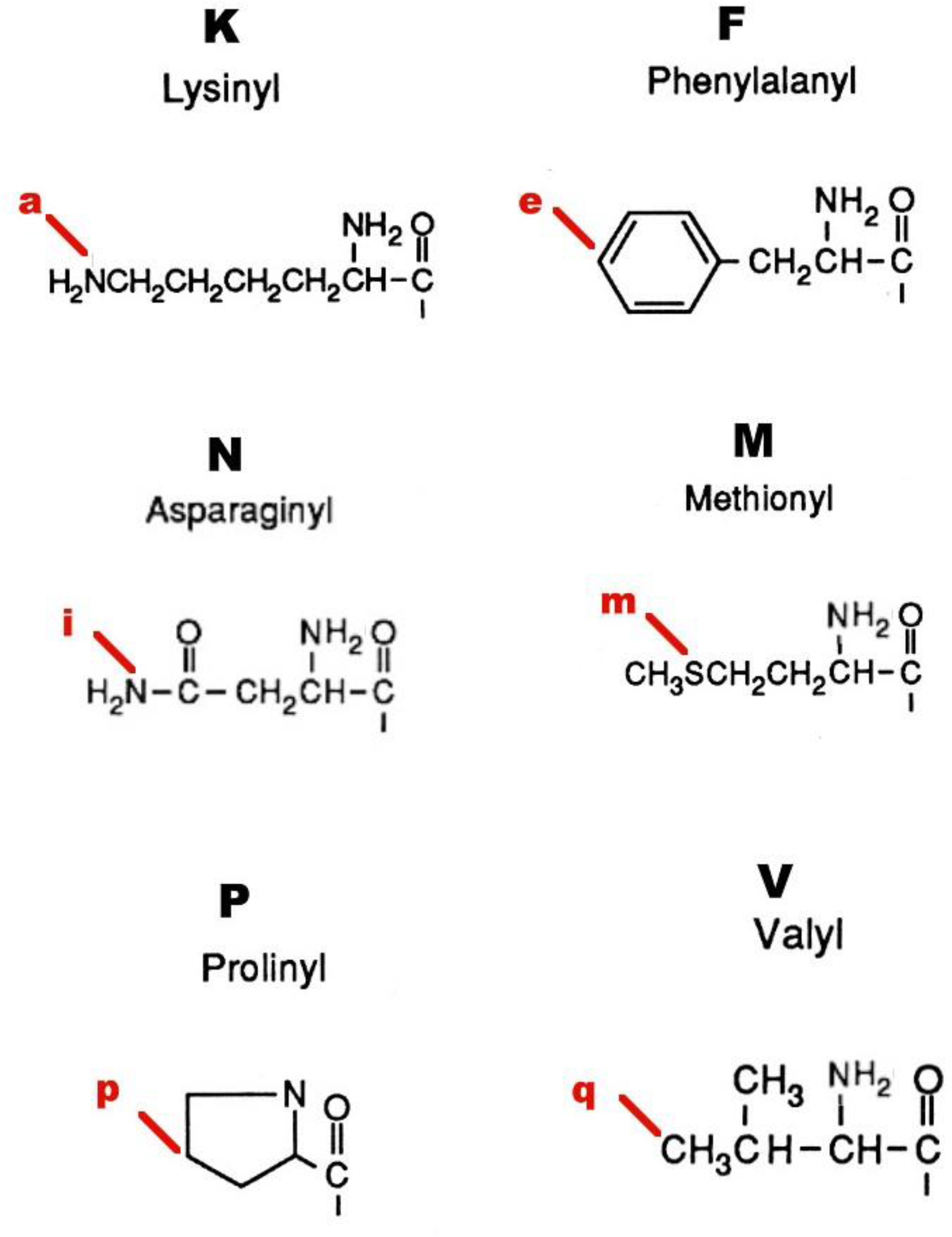
Atoms for interatomic distances.

The human equivalent of EPL143 is EPL141, with sheep V9 of sSgII-14 replaced by N9 of hSgII-14. A Del Re charge analysis shows that the C-3 point charge in these valine and asparagine residues in the peptide series is very closely similar (EPL143 = 0.293, EPL141 = 0.294, EPL001 = 0.299), suggesting that this carbon atom would be equally recognisable by both human and non-human versions of a receptor. EPL141 has much the same interatomic distances as EPL143 and a virtually identical value for *P* of 0.999͘ . These two 14mers had roughly the same inhibitory activity on prostate cancer cells in vitro (Fig. 5). For its part, EPL001 delivers an extravagant *P* value of 0.000058 because it is a different gapping of EPL143’s active face residues, without a change in their order. This apparent statistical upholding of the null hypothesis of SgII-unrelatedness is an artefact of spacing: there are 50 aa between the active face residues (15 measurements) of EPL143, **<u>M</u>**xxxxx**<u>KPVF</u>**x**<u>N</u>**xx; 72 between the equivalent residues in EPL001, **<u>MKP</u>**xxx**<u>V</u>**xxF**<u>N</u>**xx. The order of the active face residues *is* changed in the 14mer scrambled control peptides: EPL010 (e.g. xx**<u>N</u>**x**<u>VMK</u>**x**<u>FP</u>**xxxx, *P* = 0.009, s), inactive in a mouse renal cell assay in vitro (Haylor et al, 2009); EPL030 (e.g. xx**<u>KMN</u>**xxxxx**<u>PVF</u>**x, *P* = 0.024, s); & EPL040 (e.g. xx**<u>KNM</u>**xxxxx**<u>PVF</u>**x,*P* = 0.00079, s), regardless of which of the 3Ks and 2Ns are selected as the basis for measurements in the three peptides. So these disordered control sequences *are* SgII-unrelated.

A key comparison is that between the sSgII-14 benchmark EPL143 and the hexapeptide EPL601, which is the proposed sSgII related hormonal active face in contiguous form and the ovine bioactivity benchmark to boot (Fig. 5). EPL143’s active face residues, **<u>M</u>**LKTGE**<u>KPVF</u>**K**<u>N</u>**NI, are subject to a gapless compression to provide EPL601’s **<u>MKPVFN</u>**, with the 6mer not significantly dissimilar to the 14mer benchmark: *P* = 0.078. The 15 measured interatomic distances (Å) are these, ranked by similarity, with the most similar first and the least similar last, giving EPL143 first, EPL601 second and the ratio of EPL601’s distances to those in EPL143 in italics: i-q (i.e. side chain N12 for EPL143/N6 of EPL601 to side chain of V, hereafter ‘N-V’ etc), 6.08 vs 6.11 *(1.00)*; e-p (F-P), 11.36 vs 11.49 *(1.01)*; e-q (F-V), 9.19 vs 8.83 *(0.96)*; a-p (K-P), 7.91 vs 8.53 *(1.08)*; p-q (P-V), 6.81 vs 6.01 *(0.88)*; a-q (K-V), 12.25 vs 10.46 *(0.85)*; e-i (F-N), 11.54 vs 9.49 *(0.82)*; a-e (K-F), 11.88 vs 9.65 *(0.81)*; i-p (N-P), 5.96 vs 7.15 *(1.20)*; m-p (M-P), 12.63 vs 8.9 *(0.70)*; m-q (M-V), 12.79 vs 8.85 *(0.69)*; a-m (K-M), 14.20 vs 9.36 *(0.66)*; i-m (N-M), 16.43 vs 10.28 *(0.62)*; a-i (K-N), 11.98 vs 4.94 *(0.41)*; & e-m (F-M), 19.44 vs 5.37 *(0.27)*. EPL143 and EPL601 have in common the contiguous sequence **<u>KPVF</u>**. The seven biggest discrepancies in the list relate to the flanking residues M & N.

Tucking in these aa to make EPL601, by eliminating EPL143 residues posited to be uninvolved in binding (x**<u>LKTGE</u>**xxxx**<u>K</u>**x**<u>NI</u>**), potentiates activity (Fig. 5).

Why is the EPL001 scrambled control peptide EPL030 active, for example in a bone marrow cell assay (Fig. 6), more so indeed than EPL001? There is a significant probability of structural dissimilarity with the SgII benchmark, as we have seen: *P* = 0.024. This implies that the activity seen is SgII unrelated. Nonetheless, by happenstance EPL030 possesses in contiguous form part of the proposed SgII-related active face, **<u>PVF</u>**, with EPL030’s PVF measurements matching those of EPL143: e-p (F-P), EPL143’s 11.36 vs EPL030’s 10.90 *(ratio EPL030:EPL143, 0.96)*; e-q (F-V), 9.19 vs 9.31 *(1.01)*; & p-q (P-V), 6.81 vs 5.82 *(0.85)*. Besides being active in mammalian systems, EPL030 displays *anti-fecundity* activity in *C. elegans* (Davies & Hart, 2008). EPL030’s C-terminus is IEPVFT, which as a hexapeptide is EPL036. The PVF in EPL036 is a better match to the equivalent motif in the SgII benchmark EPL143 (MLKTGEK**<u>PVF</u>**KNNI), *P* = 0.995, than is EPL030’s PVF. EPL036’s PVF measurements also align with those of the PVF-containing active face hexamer EPL601: e-p (F-P) = 11.40 for EPL036 vs 11.49 for EPL601 *(ratio EPL036:EPL601, 0.99)*; e-q (F-V) = 9.34 vs 8.83 *(1.06)*; & p-q (P-V) = 7.03 vs 6.01 *(1.17)*. The possession of PVF presumably accounts for the bioactivity of EPL036 in a mammalian cell assay (Fig. 7) and for its *pro-fecundity* activity in *S. siamkayai* (Fig. 2). PVF in EPL036 is +/− in terms of tri-residue topology, the same co-accessibility as is seen for the PVF in the hexamer benchmark EPL601. Looking at the other end of EPL030, why is its N-terminal hexamer EPL630 *anti-fecundity* in *S. siamkayai*? The chi-squared analysis says that EPL630 is not significantly dissimilar to EPL143: *P* = 0.088. The active face residues in EPL630, including a reverse MK doubleton, can be represented thus: KL**KMN**G. The tri-residues KMN in EPL630 are co-accessible in the counter-clockwise orientation only, in which viewing the three side chains look like the legs of a tripod. KMN accessibility is likewise counter-clockwise only in the 14mer benchmark EPL143 but clockwise only in the 6mer benchmark EPL601. The distance measurements available for EPL630 are a-i (K3-N), a-m (K3- M) and i-m (N-M). These figures are 9.97, 10.79 & 8.62 for EPL630; 11.98, 14.20 & 16.43 for EPL143 (K7, N12); and 4.94, 9.36 & 10.28 for EPL601 (K2, N6). (For comparison, EPL630’s parent EPL030 has a K3-N of 12.39, a K3-M of 12.18 and an N-M of 7.27.) Recollecting, the active face residues are M, K, P, V, F & N. Among these K is contiguous while M and N are non-contiguous outliers (Fig. 1). Starting with K in the analysis, K7-N12 in EPL143 is 11.98, which is close to K3-N in EPL630, at 9.97, and a long way from EPL601’s K2-N6 of 4.94.

EPL601’s figure is likely to be less true to life than that of EPL143, as its F & N are unrealistically contiguous: MKPV**<u>FN</u>**. EPL143’s F & N are more realistically gapped: MLKTGEKPV**<u>F</u>**K**<u>N</u>**NI. The K-M figures are most alike between EPL630 and EPL601, at 10.79 vs 9.36. These residues are doubletons in both hexamers: KL**<u>KM</u>**NG & **<u>MK</u>**PVFN. EPL143’s figure of 16.43 reflects a five-residue gap: **<u>M</u>**LKTGE**<u>K</u>**PVFKNNI. Finally, the N-M situation is as follows: EPL630 (KLK**<u>MN</u>**G, 8.62), EPL601 (**<u>M</u>**KPVF**<u>N</u>**, 10.28) and EPL143 (**<u>M</u>**LKTGEKPVFK**<u>N</u>**NI, 16.43). The likeness is between the hexamers once more, surprisingly given MN is a doubleton in one and separated by four intervening residues in the other. Potentially accounting for the activity of EPL630 in nematode reproduction, then, are a sharing with EPL143 of stereospecificity and a key metric (K-N) and a sharing with EPL601 of the two other available metrics (K-M, N-M).

Referring again to the parent, EPL030, the sequence of this can be compared to that of EPL040, with differences underlined: KLK<u>MN</u>G<u>KN</u>IEPVFT vs KLK<u>NM</u>G<u>NK</u>IEPVFT. EPL040 is far more dissimilar in a statistical sense than EPL030 to EPL143, yet it has near identical *anti-fecundity* activity to EPL030 in *C. elegans* (Davies & Hart, 2008). The MK reversal in EPL030 and in EPL030’s N-terminal hexamer EPL630 is not present in EPL040. Apart from PVF, the only other similarity between EPL030 & EPL040 on the one hand and EPL143 on the other is the interatomic distance a-p (K-P). In fact, an a-p (K-P) correspondence is the sole commonality running through the active peptide series: EPL001 = 8.74, EPL016 = 8.30, EPL030 = 8.91 *(ratio to EPL143, 1.13; ratio to EPL601, 1.04)*, EPL040 = 9.22 *(ratios: 1.17, 1.08)*, EPL104 = 8.71, EPL120 = 8.39, EPL140 = 7.99, EPL141 = 8.06, EPL142 = 7.68, EPL143 = 7.91, EPL545 = 8.79, EPL600 = 8.57, EPL601 = 8.53 & EPL901 = 7.66. These figures all relate to contiguous **<u>KP</u>** residues except those for EPL030 and EPL040, with the arrangement in these being xx**<u>K</u>**xxxxxxx**<u>P</u>**xxx, remarkably. This suggests that the *anti-fecundity* activity of EPL030 and EPL040 is due to their possession of **<u>K</u>**xxxxxxx**<u>PVF</u>**, not just the **<u>PVF</u>** shared with their *pro-fecundity* C-terminal hexamer EPL036.

There are 11 ovine-related bioactive peptides in Table 1, when the inactive EPL801 is removed from consideration along with the human-related peptides (EPL140, EPL141, EPL600 & EPL800) and the scrambled peptides (EPL010, EPL030 & EPL040) and their progeny (EPL036 and EPL630). Each of the ovine peptides has all or some of the active face residues in what is predicted to be the endogenously correct order, though not necessarily contiguously. For each of the 11 peptides modelled up to 15 interatomic distances are available (S7). For each distance in angstroms, the range (i.e. maximum value minus minimum value) is as follows, ranked by size: a-p (**<u>K-P</u>**), range 1.13; i-q (**<u>N-V</u>**), 3.18; p-q (P-V), 4.18; e-q (**<u>F-V</u>**), 4.82; a-q (K-V), 4.93; i- m (N-M), 6.15; a-m (**<u>K-M</u>**), 6.28; e-i (**<u>F-N</u>**), 7.26; m-q (M-V), 8.46; m-p (**<u>M-P</u>**), 8.95; e-p (F-P), 9.05; i-p (N-P), 10.94; a-e (K-F), 12.92; a-i (K-N), 13.56; & e-m (F-M), 14.07. Of the 10 tightest ranges six (emphasized) are those relating to the distances *within* MKP and *within* VFN, while the five loosest ranges are *between* MKP and VFN. This is consistent with the active face aa acting as two modules of three residues each, with the distance between the residues within each module more critical than the distance between each module. The K-P commonality anchors the forward module, having the tightest range overall, with K-M only in 7^th^ position and M-P in 10^th^. V related ranges surprise by being ranked 2^nd^, 3^rd^, 4^th^, 5^th^ & 9^th^.

Unimodular tri-residue autonomy is indicated by both ends of EPL001 being independently active: **<u>MKP</u>**LTG (EPL016) and K**<u>V</u>**KE**<u>FN</u>**NI (EPL018, see Discussion). EPL590 also displays unimodular activity: AAAAAGK**<u>V</u>**KE**<u>FN</u>**NI. Activity with incomplete bimodularity is evinced by EPL901 (**<u>M</u>**LKTGE**<u>KPV</u>**, the first 9 residues of the EPL143 benchmark), EPL545 (**<u>MKP</u>**LTAAAAE**<u>FN</u>**NI) and EPL104 (**<u>MKP</u>**LTGK**<u>V</u>**KEAAAA). A full complement of bimodular aa is evinced by EPL001 (**<u>MKP</u>**LTGK**<u>V</u>**KE**<u>FN</u>**NI), EPL120 (**<u>MKP</u>**LTGK**<u>V</u>**KE**<u>FN</u>**NIKGFGVI), EPL142 (**<u>M</u>**LKTGE**<u>KPV</u>**K**<u>FN</u>**NI), EPL143 (**<u>M</u>**LKTGE**<u>KPVF</u>**K**<u>N</u>**NI) and EPL601 (**<u>MKPVFN</u>**).

EPL120 was markedly more inhibitory than EPL001 (Fig. 6). Why? Even though EPL120 is half a dozen residues longer than EPL001, there are only relatively minor differences between these peptides in terms of the 15 measured interatomic distances, with none differing by more than an angstrom. For both EPL001 and EPL120 the side chains of MKP are accessible for binding when this tri-residue is viewed in either clockwise or counter-clockwise orientations (Table 1). VFN is available for binding in both peptides when this tri-residue is in a clockwise orientation but whereas VFN is *unavailable* in EPL001 when viewed counter-clockwise, it *is* available in EPL120. Extending EPL001 to EPL120 seemingly induces a *positive* conformational change in terms of receptor binding.

The inactive EPL801 is EPL601 with EPL001’s C-terminal doubleton tacked on: MKPVFN<u>NI</u>. The addition was not predicted to reduce activity as EPL120 has NI C-terminally and 6 other residues besides. The biggest deviations in interatomic distances between EPL601 and EPL801 are these: a-i (K-N), 4.94 vs 5.87 for EPL801 *(1.19)*; i-p (N-P), 7.15 vs 9.26 *(1.30)*; & i-q (N-V), 6.11 vs 10.23 *(1.67)*. The biggest difference between the benchmark EPL143 and EPL801 is likewise i-q (N-V): 6.08 vs 10.23 for EPL801 *(1.68)*. None of the divergencies in these ‘main differences’ comparisons relate to intramodular distances within MKP, so with that module alone EPL801 might be expected to be cell inhibitory, along the lines of EPL016 (**<u>MKP</u>**LTG, Fig. 7), but it isn’t. In regard to the N-V elongation in EPL801 the relevant residues are these: MKP**<u>V</u>**F**<u>N</u>**NI. Within EPL601, the MKP of the active face hexamer’s **<u>MKP</u>**VFN is accessible only in a clockwise orientation, while its VFN is accessible only in a counter-clockwise orientation: + −/− + (Table 1). For comparison, EPL120 is + +/+ + and EPL143 is + −/+ +. The two trios in EPL601 are arranged with complementary accessibility, hinged at proline. The accessibility for EPL801’s MKP is again clockwise, but VFN is inaccessible this time in both orientations: + −/− −. Unlike in EPL601’s VFN counter-clockwise orientation, EPL801’s N6 is not co-accessible (Fig.9), instead projecting back into the complementary MKP space, providing steric hindrance. Extending EPL601 to EPL801 evidently induces a *negative* conformational change in terms of receptor binding.

**Figure 9.**
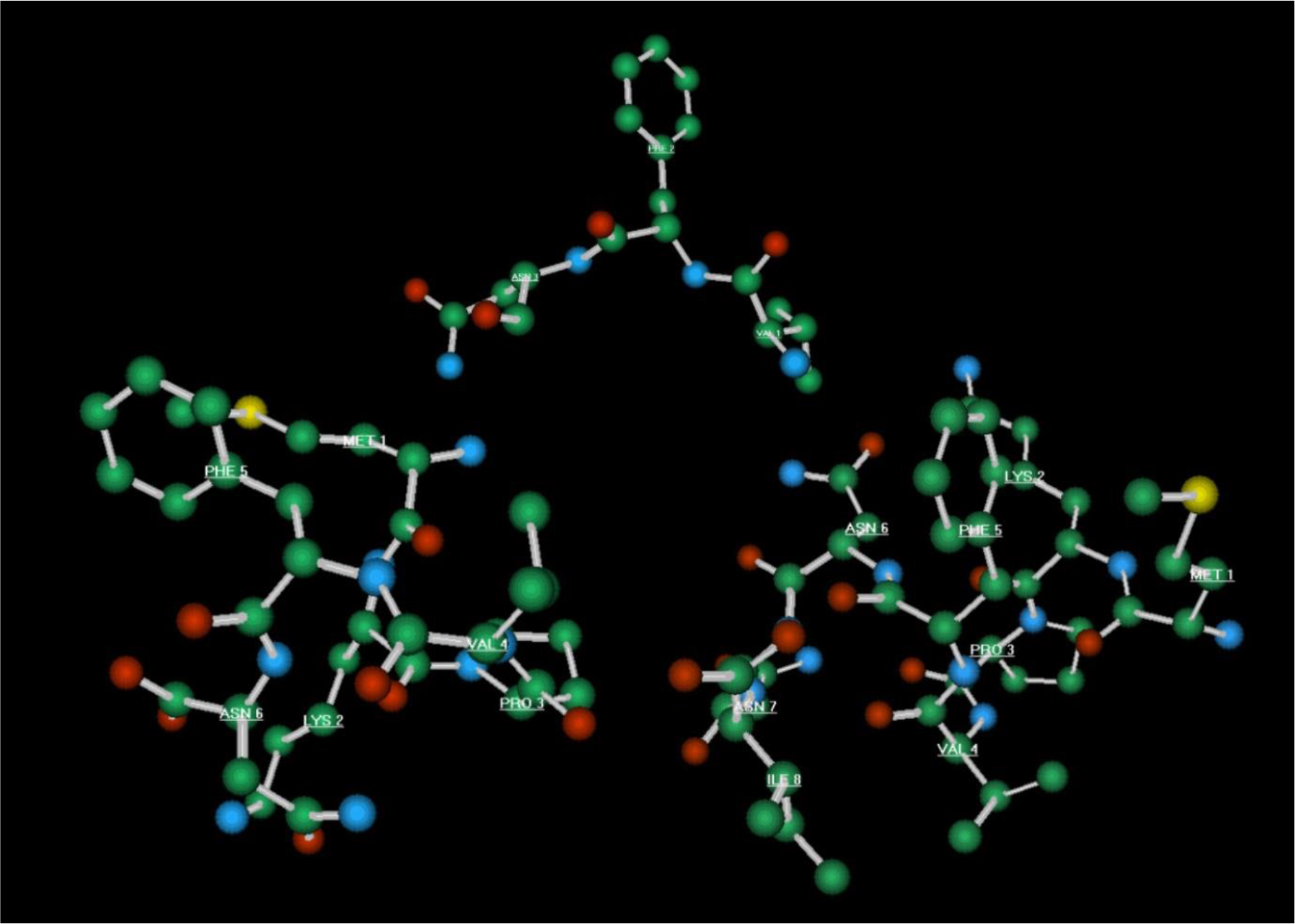
Molecular models. Minimized ball and stick representations in silico of EPL601 (MKPVFN, left) and EPL801 (MKPVFNNI, right), with the tripeptide VFN for reference (top, centre). (Hydrogen atoms deleted for clarity.) In the tri-residue topology analysis of Table 1 this is ‘VFN Θ’. The viewing starts at V (VAL 4), moving counter-clockwise to F (PHE 5) then N (ASN 6). The VFN side-chains are in the forward plane in EPL601 and accessible together for binding (+). In EPL801 they are not co-accessible (−), as ASN 6 projects rearwards.

The relative sizes of the peptides can be addressed by enclosing each of the molecular models in silico in its own tight graphical box. The dimensions of these boxes in angstroms (Length, Width, Depth) and their volume (V) in cubic angstroms are as follows, for selected peptides: **EPL001**, L 26.65 x W 17.65 x D 17.46 = V 8,212.70; **EPL120**, L 33.57 x W 16.98 x D 22.83 = V 13,013.52; **EPL143**, L 26.06 x W 19.56 x D 17.87 = V 9,108.94; & **EPL601**, L 13.87 x W 15.21 x D 14.83 = V 3,128.58. In the context of accessing a putative common receptor, the fourfold disparity in volumetric size between the bioactive peptides EPL120 and EPL601 seems especially surprising. A box able to enclose just the proposed active face residues M, K, P, V, F & N in the 14mers EPL001 and EPL143 and in the 20mer EPL120 has the dimensions L 22.57 x W 12.53 x D 16.47 = V 4,657.75. The length of this is >60% more and the volume ∼50% more than the corresponding dimensions of the box for EPL601 itself, whose sequence is MKPVFN.

## DISCUSSION

An aspiration in biology is control of fundamentals. One fundamental is reproduction. The results described here confirm that a hexapeptide of aa sequence IEPVFT (EPL036 in Table 1) is a reproductive activator, providing among *Steinernema siamkayai* entomopathogenic nematodes a superabundant monoculture of the economically important infective juveniles, while another hexamer, KLKMNG (EPL630), is revealed to be a novel reproductive inactivator. The peptides were provided at commencement ambiently via an agar substrate. The effects were dependent on dose but not on the number of worms present at the start of the study. In a study in *C. elegans*, genital tract localisation was observed after ambient exposure to a 19mer extended version of the 14mer master peptide EPL001, C-terminally labelled with fluorescein (Davies & Hart, 2008). Ingestion can be assumed to explain this uptake, rather than cuticular diffusion, and the same can be assumed here for *S. siamkayai*, but route of ingress has not been investigated and neither has the molecular mode of action. The present work is a replication study for EPL036. With two peptide dose levels and two worm population levels, there was a double duplication inherent in a study that was in any case prefigured by a pilot study conveying the same result (S3) by a new principal investigator (SM) in a new species of nematode using peptide from a different supplier.

The provenance and activity of the reproductive peptides are strange indeed. EPL036 is the tail end, ‘Back 6’, and EPL630 the ‘Front 6’ of EPL030, a 14mer peptide of sequence **<u>KLKMNG</u>**KN**<u>IEPVFT</u>**. EPL030 was synthesized for use in experiments as a control compound. The EPL030 sequence is a random scrambling of the master test sequence to emerge from a hormone discovery project, MKPLTGKVKEFNNI, synthesized as EPL001. Why do disparate terminal fragments of a scrambled-sequence control peptide show any bioactivity at all? Divergent activities indicate that non-specific effects are probably not in play. Agonism and antagonism involving a single biological system might be involved or alterations in different systems. EPL030’s Back 6, EPL036, is *pro-fecundity* in the nematode *Caenorhabditis elegans* (and in guppy fish and frogs among other organisms) and is likewise *pro-fecundity* here in *S. siamkayai*, while its parent EPL030 is *anti-fecundity* in *C. elegans*, dose dependently. In line with the latter observation, EPL030’s Front 6, EPL630, is *anti-fecundity* in *S. siamkayai*, while as yet untried in *C. elegans*. Meanwhile, the grandparental EPL001 14mer in *C. elegans* is *pro-fecundity* (Davies & Hart, 2008; Davies et al, 2015). How is all this to be untangled? EPL001 is a found sequence, as described in the Introduction. As a 14mer synthetic peptide it has been represented elsewhere as a minimized ball and stick model in silico (Howlett et al, 2019). EPL001 amounts to a 20% N-terminal fragment of Candidate 7500, an approximately 70 aa candidate polypeptide from gel electrophoresis and MS for a postulated inhibitory hormone, yet EPL001 displays on its own in vivo the two crucial features of the sought-for hormone: anti-organotrophic activity (dose-dependently) and reproductive modulation. An anti-EPL001 antibody administered alone potentiated compensatory renal growth in the rat after unilateral nephrectomy (Haylor et al, 2009). This is consistent with immunoneutralisation of an EPL001 related endogenous inhibitor of kidney mass. Either EPL001 is a true product of the hormone hunt or its presence is an exceedingly bizarre coincidence. An attempt has been made here to eliminate a mild, enduring antiproliferative effect of EPL001 on human mesenchymal bone marrow stem cells in culture, via alanine substitution of substrate-tethered peptides. Three EPL001-related 14mer peptides were tried, having alanines in positions 1-5 (EPL590), 6-9 (EPL545) and 11-14 (EPL104), together with EPL001 itself. All were associated with 23-37% fewer cells then controls after 14 days, with unchanged stem cell phenotype and without apparent toxicity. If the active site were in the centre section of EPL001 then the 6-9 alanine substitution should have knocked it out, but it didn’t. With all the peptides being antiproliferative this suggests that both ends of EPL001 are active, independent of one another; an odd conclusion, that forestalled the use of an alanine ladder. EPL001 was inhibitory in this system regardless of which end of the peptide was tethered to the substrate, though the C- terminally tethered item was the more active of the two. This suggests that freed of steric hindrance the main activity is at the N terminus. Yet in a study in vitro involving human dental pulp cells – a more reliable testbed for inhibitory peptides than bone marrow stem cells (Rogers, 2018) – the EPL001 Front 6 and Back 8 peptides and the master peptide itself at 5µM (single starting dose) in the cell medium were mildly antiproliferative in the order MKPLTG (EPL016)<MKPLTGKVKEFNNI (EPL001)<KVKEFNNI (EPL018). The last-mentioned reduced cell numbers by ∼17% over 7 days (Hart, 2008). In the present work the EPL001 Front 6 peptide MKPLTG (EPL016) at 1µM in the medium reduced the population of rat bone marrow cells in culture by 18% in 24h, with peptide-untreated controls increasing in numbers by about the same percentage (Fig. 7). (The reproductive activator EPL036 was cytostatic.) In a prior assay using the same system, cells exposed to EPL001 were significantly fewer than controls but there was no absolute reduction in numbers, as there was via apoptosis with the EPL001 control peptide EPL030 and with EPL120, an extended 20mer version of EPL001 (Fig. 6). EPL001 is sometimes devoid of inhibitory activity in vitro altogether (Fig. 5; Haylor et al, 2009) and generally unpredictable (Rogers, 2018), while EPL120 has mostly proved more cell-inhibitory than EPL001 in the experimenters’ hands. For example, EPL120 showed dose-dependent inhibition of the (exogenous) IGF-1 stimulated growth in cell numbers of MCF 7 human breast cancer cells over 72h across the EPL120 dose range 0.04 nM - 4 µM (Hart, 2008). Similar results have been obtained with EPL001, albeit evoked more variably, yet the 14mer was without significant antitumour effect in the MCF-7 xenograft nude mouse model. That EPL001 ‘works’ in vivo is demonstrated by inhibition of compensatory renal growth in the rat (Haylor et al, 2009). The renal study failed to deliver an in vitro correlate. It was considered that the in-life result might involve an indirect, more ‘endocrine’ disruption in what is an IGF-1 related model (Haylor et al, 2000).

Returning to EPL030, the scrambled-sequence EPL001 comparator, this has engendered perplexity by displaying the same activity as EPL001 across multiple mammalian systems. Thus, EPL030 suppressed human prostate cancer cells in vitro more than EPL001 in two laboratories and the proliferation of rat bone marrow cells in vitro in a third. Infused into the third ventricle of sheep, EPL030 was associated with the same pattern of anterior pituitary hormone changes in the peripheral blood as EPL001, but a GH elevation which eluded statistical significance for EPL001 (+25.4%, *P* = 0.194) reached it for EPL030 (+43.8%, *P* = 0.039). The latter observations are consonant with others: hormone production from pituitary cells in vitro is altered in the same direction by exposure to EPL001 and EPL030 (Hart, 2008; S4); ovine hypothalamic immunostaining has been demonstrated with an anti-EPL001 antibody (see Introduction), without pituitary staining (Hart et al, 2017); an aqueous extract of rat hypothalamus had an antiproliferative effect on rat bone marrow cells in vitro, except when subject to prior immunodepletion with an anti-EPL001 antibody, unless EPL001 was added as well during the immunodepletion process, achieving preabsorption (Hart et al, 2017). At the highest dose in maximum tolerated dose studies in mice, EPL030 caused more inappetence and consequent weight loss than did EPL001, within a picture of low toxicity for both peptides. In a mouse xenograft study the average final tumour mass was 20% lower (ns) for EPL030 than the values for EPL001 or PBS-only controls, which two latter values were similar at termination (S5 Fig. 3). Significantly slower tumour growth might be elicited by a higher dose of EPL030 (or EPL001) or by a switch to more promising effectors (EPL120, EPL600, EPL601). Note that in these mammalian studies EPL030 had the same activity direction as EPL001, only more so. In contrast, as we have seen, EPL030 is *anti-fecundity* in *C. elegans*, as is the closely similar control peptide EPL040, while EPL001 is *pro-fecundity*. What does it mean when a control compound gains a life of its own and behaves like the test compound in mammalian systems or oppositely in a species of nematode? When the test compound itself, EPL001, is enigmatic, perplexity thickens. Stochastic non-specific effects or is there illumination to be had in terms of structure-activity relationships? As between EP001 and EPL030, Spearman’s rank correlation is 0.30, a weak positive correlation betokening a degree of randomisation for EPL030 (S2).

Peptide sequence scrambling may involve just an active motif (Willman et al, 2010) or the entire length of a peptide (Xia et al, 2020). Scrambling methods are seldom described. Control peptides usually oblige with inactivity.

Peptide studies frequently in fact involve generating a scrambled-sequence (anagrammatical) peptide – that is, a control peptide of the same aa composition and MW but devoid of relevant sequence-related activity, though possibly confounding for other reasons (e.g. nutrient effect, fortuitous assembly of an unrelated biological motif etc.). This is the provision of a negative control (Moser, 2020). How best to scramble aa sequences? *Rule-based randomisation* is suggested, rather than unregulated randomisation used here. Accept each random ‘pick’ only if the aa differs in position from the corresponding residue in the source peptide and, in the case of all residues save the first, accept the pick only if the preceding pick is a non-neighbour in the source peptide. In the example of EPL001, MKPLTGKVKEFNNI, the first pick to be accepted can be anything except M; the second pick is any of the remaining 13 residues except one of the three Ks or a neighbour within EPL001 of the first pick; the third pick is anything left except P or a neighbour of the second pick; and so on. If the last pick is I, start again. No three aa should be in the same order across the whole sequence, regardless of gapping. (Additional rules to achieve a ‘most different’ combination could include non-acceptance of residue picks of a similar type, e.g. acidic, basic, hydrophobic, and acceptance only on the basis of dissimilar gaps between non-contiguous residues.) With hindsight, then, the technique employed here of ‘aa residues pulled randomly from a hat’ should have been accompanied by the rule ‘reject residue if it occupies the same position in the source peptide or if the prior pick is a neighbour in the source peptide or if tri-residue disorder has not been achieved across the entire sequence’. There are three fortuitous positional matches between EPL001 and EPL030: MKPLT**<u>GK</u>**VK**<u>E</u>**FNNI and KLKMN**<u>GK</u>**NI**<u>E</u>**PVFT. Six EPL001 residues ended up in EPL030 with the same neighbours: **<u>M</u>** with **<u>K</u>**; **<u>G</u>** with **<u>K</u>**; and **<u>N</u>** with **<u>I</u>**, as in **<u>MK</u>**PLT**<u>GK</u>**VKEFN**<u>NI</u>** and KL**<u>KM</u>**N**<u>GK NI</u>**EPVFT. Referring to the proposed active face residues, **<u>MKP</u>** is present in that tri-residue order, variously spaced, in all three control peptides – EPL010, EPL030 and EPL040 – and **<u>V</u>·<u>F</u>·<u>N</u>** is to be found additionally in EPL010 (Table 1).

A failure of randomisation is available for inspection in the form of EPL630. This hexamer of sequence KLKMNG was anticipated to be an inactive control in nematode breeding studies, but turned out to be a reproductive inactivator. The KM in EPL630 is a reversal of the MK in master test peptide EPL001 (**<u>MK</u>**PLTGKVKEFNNI). Both M and K are among the sSgII-70 proposed active face residues – M, K, P, V, F & N – which were so designated in the Introduction on the basis of *correlated prominence* in terms of evolutionary conservation (MxxxxxK, PV), purification prevalence (all), previously ascertained bioactivity (P in position 3), preliminary computational modelling (FN) and sequence gridding (Fig. 1’s grid-topping MKP, plus VFN). The correspondence in full of EPL630 to the proposed hormonal active face is KL**KMN**G. So, why is EPL630 reproductively suppressive in a species of nematode? *Tri-residue topological conformity* (in this case with EPL143), defined as close correspondence in terms of stereospecific space occupancy and interatomic separation of identical trios of aa side chains within any designated pair of peptides, whether the aa are in the same sequence order or not. The KMN side chains are arranged like a splayed tripod, suggestive of binding capability, and they are a stereospecific match for the corresponding residue side chains in the 14mer benchmark EPL143: **<u>M</u>**LKTGE**<u>K</u>**PVFK**<u>N</u>**NI (= sSgII-14 of Fig. 1). EPL630 and EPL143 are thus in topological conformity with each other, with EPL630’s trio of aa side chains and EPL143’s displaying superimposable co-accessibility, i.e. same-handedness. This is not true of EPL630 and the 6mer benchmark EPL601, where the trios show opposite-handed co- accessibility. Another likeness with EPL143 is the gap between two of the legs of the tripod, the K-N distance. This is a key metric as while K is posited to be one of five active face residues (MKPVF) from the amino terminus of sSgII-70, and contiguous at that (i.e. K is locatable in space as covalently bound to P, Fig. 1), N is the sole residue from its carboxy terminus. Joining N to F as in EPL601 (MKPV**<u>FN</u>**) is unlikely to be true to life in terms of the separation of these non-contiguous residues, whereas the **<u>F</u>**x**<u>N</u>** gapping of EPL143 is anticipated to provide a more accurate K-N metric. The distances between the other legs of the tripod, K-M & N-M, are more in line with those in EPL601. A bimodularity concept arose out of the analysis of interatomic distances. EPL630’s KMN, a tight trimer within KL<u>KMN</u>G, paradoxically displays *wide* half-complement bimodularity: **<u>MK</u>**P-VF**<u>N</u>**. Its EPL030 sibling and opposite number in reproductive terms, EPL036, IEPVFT, displays *narrow* half-complement bimodularity: MK**<u>P</u>**-**<u>VF</u>**N. These hexamers are active because the two ends of their 14mer parent EPL030 each contain three of the six active face residues: EPL630 has K, M & N, non-contiguous one with another in the active face but contiguous in EPL630; while EPL036 has the easier to analyse P, V & F, contiguous in the active face *and* in EPL036. The KMN distances in EPL630 (a-i, a-m, i-m) and its parent EPL030 are different, so in this respect neither throws light on the other’s bioactivity. This is a challenge as both have *anti-fecundity* activity in nematodes, EPL630 in *S. siamkayai* here, EPL030 in *C. elegans* previously. EPL030 matches the proposed active face in this fashion: KL**KMN**GKNIE**<u>PVF</u>**T. The EPL030 analysis in the molecular modelling identified a four-residue motif as potentially important in a bioactivity assessment: **<u>K</u>**xxxxxxx**<u>PVF</u>**. This does not represent a failure of randomisation, as EPL001 does not have a K in position three or a PVF anywhere. Instead, at issue is the fortuitous assembly of a partial proxy of the proposed sSgII-related active face, M**·<u>KPV</u>·<u>F</u>·**N, the whole of which in contiguous form is EPL601. On this analysis EPL030 is a crypto-test compound because of its possession not just of **<u>PVF</u>** but of **<u>K</u>**xxxxxxx**<u>PVF</u>**. The K-P (a-p) correspondence of EPL030 (and of the closely similar EPL040) with those in EPL601 and EPL143 is especially surprising given that in both the benchmarks **<u>KP</u>** appears as a doubleton. The C-terminus of EPL030 yields EPL036, IEPVFT. While EPL030 is *anti-fecundity* in *C. elegans*, EPL036 is contrarily *pro-fecundity* in *C. elegans* and in *siamkayai*. EPL036 has only the **<u>PVF</u>** of its parent’s **<u>K</u>**xxxxxxx**<u>PVF</u>**. EPL036 has the same co-accessibility for PVF binding as the hexamer benchmark EPL601, +/−. The hormone hypothesis is of a conserved endocrine factor that curbs tissue overgrowth and restricts reproduction (Hart, 2014). Accordingly, the *anti-fecundity* activity of the 14mer EPL030 and its Front 6 EPL630 is in line with theory and are examples of agonism (the mimicking of an endogenous inhibitor). In this scheme EPL036 acts as a binding-accessible (+/−), narrowly bimodular (xx**<u>P</u>**-**<u>VF</u>**x) antagonist, as per a prediction of antagonistic activity from prior molecular modelling (Davies et al, 2015), to counteract an endogenous inhibitory hormone. EPL036 disinhibits reproduction by blocking a default hormonal OFF signal that EPL630 mimics.

The aa sequence of EPL001 can now be deconvoluted. The sSgII related proposed active face **<u>M</u>·<u>KPV</u>·<u>F</u>·<u>N</u>** is found in ovine-derived EPL001 dispersed thus: **<u>MKP</u>**LTGK**<u>V</u>**KE**<u>FN</u>**NI. This renders comprehensible the ‘both ends active’ results of alanine substitution by sector, assuming that less than a full complement of aa is sufficient for activity. Each of the three alanine-substituted EPL001 peptides has at least half of the six active face residues, explaining retention of antiproliferative activity within the bimodularity concept. The minimum necessary for stereospecificity is three points of attachment (Easson & Stedman, 1933; Ogston, 1948).

Compared with EPL601 (MKPVFN), EPL001 has a similar **<u>MKP</u>** and independently a similar **<u>V</u>·<u>FN</u>**, the distances between these two modules in the 14mer bearing no relationship to those in the 6mer. In the peptide immobilisation study in vitro, N-terminally immobilised EPL001 was less inhibitory of stem cell proliferation than C-terminally tethered EPL001. The probable reason for this is that the two effector modules have differential binding affinities to cell membrane receptors in this system, with the forward module **<u>MKP</u>** being more avid than the rearward module **<u>V</u>·<u>FN</u>**. Bimodularity as between **<u>MKP</u>** and **<u>VFN</u>** is an implicit prediction of the Fig. 1 sequence grid: in spite of P and V being covalently linked in the sSgII scheme they are in different sectors of the grid. EPL601, **<u>MKPVFN</u>**, is posited to be a biomodular effector hinged at P. The EPL601 both-modules-together binding is compromised by elongating the hexamer to provide EPL801: MKPVFN<u>NI</u>. EPL801’s N6 departs the VFN space (Fig. 9) and intrudes into the MKP space, providing steric hindrance. EPL120, the 20mer extended form of the 14mer EPL001, and EPL601, the 6mer minimisation of EPL001, are both more anti- proliferative than EPL001 in mammalian cell assays, as is the 14mer sSgII benchmark EPL143. In the tri-residue topological analysis only three ovine test peptides have the stereospecific signature ‘MKP clockwise, VFN counter-clockwise’: EPL120, EPL143 and EPL601 (Table 1). As between its two ligand modules, the unpredictable master test peptide EPL001 has adverse topology, ‘MKP clockwise, VFN clockwise’, while the inactive EPL801 merely boasts ‘MKP clockwise’. At issue with these ligands is simultaneous accessibility for binding, or the lack thereof, of two trios of aa side chains.

A graphical box enclosing a model in silico of the 6mer EPL601’s MKPVFN is six-tenths as long as a box enclosing the MKPVFN residues in the 14mers EPL001 and EPL143 and the 20mer EPL120. Ligand binding can induce changes in the 3D structure of a receptor but such conformational alterations would need to be marked to accommodate such a dimensional discrepancy. The concept of a unitary receptor for disparately sized effectors can be rescued by the same concept that applies to the ligands themselves, stereospecific bimodularity – only, in the case of the receptor, doubled up (at least). A dimeric receptor would have four stereocentres, two in each site, matching the two in each ligand molecule. In this model, EPL601’s MKPVFN would bind with the receptor’s attachment points M*’*K*’*P*’*V*’*F*’*N*’*, while a second molecule of EPL601 would be free to bind with the receptor’s dimeric attachment points, M*”*K*”*P*”*V*”*F*”*N*”*. Dual occupancy could be necessary for full receptor activation. The outsized EPL143 and EPL120 can cross-bind to M*’*K*’*P*’* & V*”*F*”*N*”* or to M*”*K*”*P*”* & V*’*F*’*N*’*, with dual occupancy again a possibility if the two receptor sites are antiparallel, M*’*K*’*P*’*V*’*F*’*N*’* vs N*”*F*”*V*”*P*”*K*”*M*”*, as shown in Fig. 10. If the receptor sites run in the same direction, M*’*K*’*P*’*V*’*F*’*N*’* vs M*”*K*”*P*”*V*”*F*”*N*”*, then cross-binding is still a possibility but not simultaneous dual occupancy, as the first molecule to bind would impede the second. The hexameric EPL601 is too short for cross-site binding, in either the antiparallel or parallel arrangements, and the same is true of the other short peptides. Dual occupancy is unrestricted, as binding within one receptor site does not preclude binding to the other. The EPL001 fragment EPL016 (**<u>MKP</u>**LTG) can bind to M*’*K*’*P*’* & M*”*K*”*P*”*, while the other EPL001 fragment, EPL018 (K**<u>V</u>**KE**<u>FN</u>**NI), can bind to V*’*F*’*N*’* & V*”*F*”*N*”*. These bindings block the other sterocentre within each receptor site but without steric hindrance of the other site’s two stereocentres. Bimodular effectors are the reproductive inactivator EPL630 (KL**KM<u>N</u>**G), which would bind to M*’*K*’*N*’* & M*”*K*”*N*”* and the reproductive activator EPL036 (IE**<u>PVF</u>**T), which would bind to P*’*V*’*F*’* & P*”*V*”*F*”*. EPL001, outsized like EPL143 and EPL120 but this time with adverse topology, can bind to any available single stereocentre in the two receptor sites: M*’*K*’*P*’*, V*’*F*’*N*’*, M*”*K*”*P*”* or V*”*F*”*N*”*. The binding to a single stereocentre might impinge on other EPL001 bindings through steric hindrance, prejudicing dual occupancy, accounting for the unpredictable activity of EPL001.

**Figure 10.**
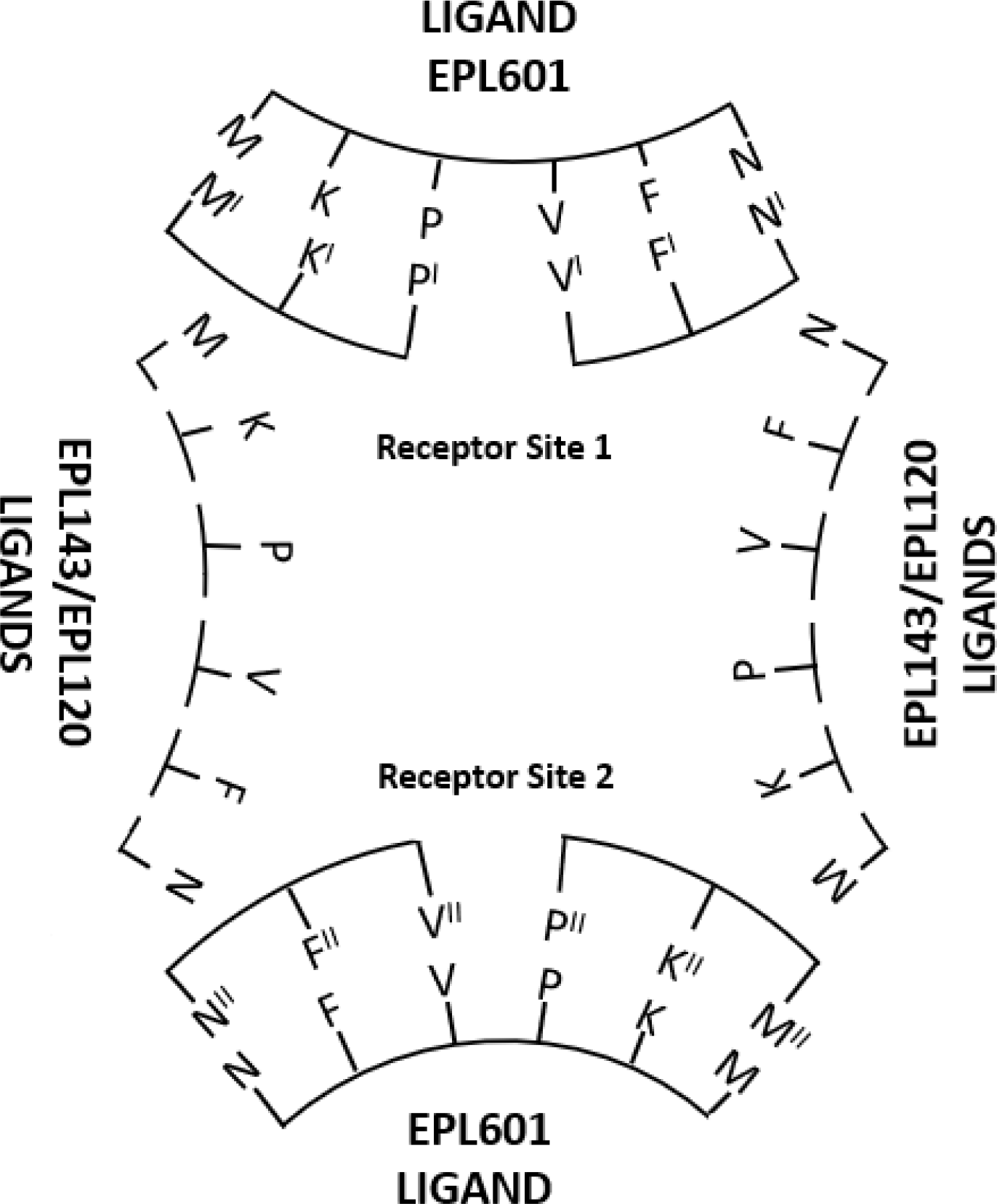
Dimeric receptor conceptualisation. A 6mer peptide (EPL601), a 14mer (EPL143) and a 20mer (EPL120) can be brought into a single interpretative scheme of bioactivity by positing a receptor comprising (at least) two binding sites, in antiparallel orientation, each site having two tri-residue modules. Ligand bimodular residues (MKP & VFN) actual, receptor residues (with superscripts) hypothetical.

The EPL030 control peptide would bind to two stereocentres within a single receptor site, as it boasts a bimodular 2/3^rd^ residue complement of **<u>K</u>·<u>PVF</u>** rather than either of EPL001’s unimodular half complements, **<u>MKP</u>** and **<u>V</u>·<u>FN</u>**. A second EPL030 molecule might well be able to bind with the other receptor site. These differences explain the bizarre finding that randomising EPL001 yields, in EPL030, a stronger agonist cell inhibitor in mammalian systems. EPL030 was deduced to be acting to *anti-fecundity* effect in nematodes as an <u>agonist</u>.

Within the inhibitory hormone hypothesis, this peptide, appreciated now as a crypto-test compound, would be expected to be an agonist inhibitor in mammalian cell assays – and it is. Why is EPL001 also inhibitory in these systems when it is a *pro-fecundity* <u>antagonist</u> in nematodes, blocking an endogenous brake? (This question is also relevant to the *pro-fecundity* peptides EPL016 and EPL036.) This is presumably because mammalian cell cultures lack the endogenous hormonal inhibitor, so EPL001 behaves as a weak agonist to inhibitory effect (ditto EPL016, **<u>MKP</u>**LTG, and EPL036, IE**<u>PVF</u>**T). The foregoing analysis supports the concept that for the peptides under review a single mode of action is at issue via a common evolutionarily conserved receptor, potentially an antiparallel dimer requiring dual occupancy for full activation.

As to receptor location, a hint is available from nematode histology and pharmacology. Concentration in the genital tract of *C. elegans* was seen after exposure to a fluorescent analogue of EPL001 (antagonist, *pro-fecundity*) but not with an analogue of EPL030 (agonist, *anti-fecundity*), from which an uninformative generalised picture was obtained (Davies & Hart, 2008).

The suggestion has been made that the body has a reproductive hormonal brake against tissue overgrowth, which brake is lifted to initiate puberty and the assumption of adult stature and whose action wanes with age, leading to a rise in cancers and the enlargement of the prostate (Hart, 2014). The factor hunt that arose from this suggestion has progressed as follows: **Inhibitory Hormone Hypothesis** ð **Candidate 7500** ð **EPL001** ð **SgII Relatedness**. The final step is supported by neuroendocrine IHC, together with MW and other circumstantial evidence of granin identity (see Introduction); an anti-EPL001 antibody immunoprecipitation leading to mammalian ‘likely SgII relatedness’ and a fruit fly homologue of SgII; unconventional bioinformatics (the ovine NNI-ome) identifying the potential relevance to EPL001 of the second sorting domain of SgII, consideration of which opened up the possibility of SgII-based epitope and active face analyses and led to the sequence grid of Fig. 1; and the 14mer constructs EPL141 and EPL143 (representing h/sSgII-9 + EPL001’s F, K & NNI) being more cell-inhibitory than straight EPL001 itself, while sSgII-9 (MLKTGEKPV = EPL901) on its own is also active (Fig. 5). Gene expression has been studied in human bone marrow stem cells in vitro using RT-qPCR (Rogers, 2018). This is in regard to the core circadian clock genes *Bmal1*, *Per2* & *Rev-ErbA* and the differentiation marker genes *OCN*, *Sox9* & *PPARy*. The hybrid 14mer EPL140 (= hSgII-9 + EPL001’s K & FNNI) induced the same pattern of changes as EPL001, with *Bmal1* upregulated amidst broad downregulation. A similar pattern was seen in human dental pulp stem cells, except that *Per2* was elevated by both 14mers as well as *Bmal1* (S6). These results are in line with both the SgII-relatedness of EPL001 and the inhibitory hormone hypothesis.

An F and a K remain unaccounted for to make up sSgII-14 (= EPL143), the shuffled version of EPL001 in Fig. 1. It is the case that there is in the UniProt sequence for sSgII (W5QEU8) this doubleton: 178**<u>FK</u>**179. Along the string there is 236**<u>NNI</u>**238, the motif deemed crucial for full binding of the anti-EPL001 antibody when exposed C-terminally. Epitope mapping gave rise to the deduction that the sought-for sSgII proteoform was the product of reverse peptide splicing within a single SgII molecule or splicing between two separate SgII molecules (Howlett et al, 2019). In the relevant location of intracellular secretory vesicles, another granin protein, chromogranin A, has been reported to be involved with insulin in an exotic form of peptide fusion splicing (Baker et al, 2018). Focusing on a monomolecular provenance and bearing in mind that r*everse splicing* involves two sequence modules being joined in the opposite order in which they appear in the parent protein, it can now be contended that a 9+61 reverse peptide splicing gives rise to ‘sSgII-70’ (pronounced ‘sheep sig two seventy’) as follows, with EPL001’s 14 aa emphasized and dots representing proposed epitope residues:

1MLKTGEKPV**<u>FK</u>**RTNEMVEEQYTPQNLATLESVFQELGKLTGPNNQKHERADEEQKLYTDDEDDIYKA**<u>NNI</u>**70

Outside the framework of the present paper, support for the sSgII-70 9+61 sequence contention is provided by mass spectrometry, with and without trypsinisation (Hart, 2021). Within the framework of the present paper, joining the V to the untethered F in Fig. 1 and the F to the untethered K (dotted lines) means that the sSgII-14 mountain range now has four peaks (M1, T4, K7 and K11), to match the four ski slopes of EPL001’s grand slalom (MKP, LTG, KVK and EFNNI). There is an amphipathic pattern of charged residues (Ks & E) between hydrophobic residues (shaded), with the Beale 4’s **<u>PVF</u>** (see Introduction) representing a hydrophobic cluster: MLKTGEK**PVF**K. Adding FK to sSgII-9 means that the quadripartite active face **<u>M</u>·<u>KPV</u>·<u>F</u>·<u>N</u>** becomes the triply non-contiguous **<u>M</u>·<u>KPVF</u>·<u>N</u>** of the heading to Table 1, Column 2. The grid predicts that after sSgII-14’s K11, the string loops off and comes back as a C-terminal NNI. This would facilitate the involvement of both ends of sSgII-70 in (i) aberrant Edman sequencing, (ii) antibody binding and (iii) the assembly of a non-contiguous active hormonal face for receptor binding. Explained by this hairpin structure is why EPL001, representing just 20% of Candidate 7500, has the activity of the sought-for inhibitory hormone. EPL001 is the aberrant reading of the two ends of the hairpin, which together contribute to the active face. Cross-linking can be suspected, as effecting the hairpin and bringing FK and NNI together, though the absence in sSgII-70 of cysteine residues (a secretogranin feature), to provide stabilising disulphide bonds, makes the chemical character of this conjectural. The EPL001 sequencing zigzag in Fig. 1 suggests the reading of a spiral: so, a cross-linked, spiralised polypeptide hairpin – with C-terminal polyanionic clustering for good measure: 48ExxDEExxxxxDDEDD63.

The Edman reagent, phenyl isothiocyanate, reacts with the α-amino groups of peptide molecules anchored to a solid phase, to liberate a cyclic derivative of the N-terminal residue for chromatographic identification (Smith, 2001). The process then repeats itself. The assumption here is that in automated sequencing the Edman reagent behaved faithfully in latching on to α- amines and that the non-provision of a database-meaningful N-terminal sequence is due to target molecule chemistry. Nothing like Edman Nonsequentialism, as it might be called, has been reported since the method was first described (Edman, 1950). Yet Edman Nonsequentialism is *not* a conjecture, it must be emphasized, but an observation, given the variant sequences obtained from the physicochemical purification campaign. The available seven sequences have much in common with one another in terms of residue incidence and position, an interpretation backed up statistically by a residue distribution analysis (S1). What is being said here is that EPL001 is not a true-to-life sequence of endogenous residues at all; but a machine artefact. The deduction is that a non-linear sequence has been obtained (except for a correct initial methionine, indicating that the target peptide is not N-terminally blocked or lacking an α-amine through being cyclic) because of the aberrant availability of free amine groups due to out-of-order peptide bond cleavages amounting to a multistage depolymerisation. The readings are ‘picked’ in (spiral) surface accessible order, MKP, LTG, KVK and EFNNI (Fig. 1), such that EPL001 encodes the source molecule’s spatial configuration, explaining why an anti-EPL001 antibody can find the SgII-related source molecule. Other factors influencing residue selection in the sequencing include steric hindrance, the presence of charged lysines N-terminally and deformation of the spiralised, looped-back, C-terminally polyanionic target molecule on the supporting membrane. The EPL001 ‘route’ taken by the sequencing has been analysed in detail (S8, Edman Nonsequentialism). The testable prediction is that putting wild-type sSgII-70 through an Edman machine would generate the EPL001 sequence or something similar.

The sequence grid of Fig. 1 is multiply informative: the active face tumbles down from the peaks represented by the evolutionarily conserved **<u>M</u>**xxxxx**<u>K</u>**; bimodularity is hinted at as between **<u>MKP</u>** and **<u>VFN</u>**; epitope residues fringe the foot of the mountain; the uninvolvement of L, T and G in active face and epitope considerations correlates with obscured positioning; the sSgII-14 track resembles the model in silico for sSgII-9 (MLKTGEKPV = EPL901); covalent bonds can be suggested in the form of V-F and F-K; the two clean zigzag tracks, one vertical for sSgII-14, one horizontal for EPL001, connote two routes around a spiral tertiary structure (S8, Fig. 3); together on the righthand side of the grid are the Beale 4 residues (PVFN), the first three of which are contiguous in sSgII-70 yet gapped in EPL001 (xx**<u>P</u>**xxxx**<u>V</u>**xx**<u>F</u>N**xx), suggesting picks around an amphipathic spiral, while the last two, FN, imply physical proximity; and there is the speculation of cross-linkage to explain the structural hairpin presumed necessary to bring together FK and NNI.

Immunoprecipitation in the inhibitory hormone project delivered SgII relatedness (Hart et al, 2017); bioinformatic NNI-omics led to the second sorting domain of sSgII (Howlett et al, 2019); whence the anagrammatical sequence grid of Fig. 1, linking EPL001 and sSgII-14, for which there would appear to be no literature precedent. In fact, though, any two anagrams can be arrayed on a grid to provide a Fig. 1 style arrangement. Consider NIGHT and THING. Range the numbers 1-5 across the top and side of a grid and express the anagrams with subscript numbers for positioning: N1I2G3H4T5 and T1H2I3N4G5. The N of NIGHT goes in column 1, row 4; I in column 2, row 3 and so on. NIGHT reads across column by column and THING reads down row by row. The letters of NIGHT can be joined up to provide one track and those of THING to provide another. But the two lines make no particular pattern together and no fresh insights are garnered by this exercise. The situation is not changed if the anagrams are *cognate*, that is, relate to one another in some way, as in the case of LISTEN and SILENT. EPL001 and sSgII-14 are obviously anagrams (the former solved to reveal the latter as the *subject*) and cognate, non-obviously (both SgII related, it is argued here), but an additional factor is key besides the provision of the right letters for an anagram: Fig.1 is a two-dimensional representation of a pair of aa anagrams in 3D space. EPL001 is a spatial mapping of sSgII-14. This is why the sequence grid is imbued with meaning, providing the contoured functional landscape referred to in the Introduction and explaining why an anti-EPL001 antibody is able to see a SgII related antigen, an otherwise unfathomable observation.

The ‘61’ part of the 9+61 reverse peptide splicing deduction is an uncomfortable one, as from the same 61-residue stretch in rat SgII is hewn the 33mer active peptide secretoneurin (SN) in its entirely and part of another active peptide, the 66mer EM66. (The equivalent sequences to rat SN and EM66 in sheep SgII are T181-Q213 and E216-M281.) From rat 61’s commencing 181FKR183 is cleaved away the dibasic site KR, by prohormone convertases, which also cut away a KR between SN and EM66. This latter KR is 214KH215 in sSgII (46KH47 in sSgII-70 numbering), rendering moot the proteolytic situation in the sheep. Of the proposed active face residues, MKPVFN, none is in rat SN and only the N of **<u>N</u>**NI is in EM66, with NNI internalised in the latter and so less visible to the anti-EPL001 antibody. A different proteolytic pathway would be involved in the production of SgII-70; different and more elaborate. Complex intracellular cascades involving post-translational modification have been described for Notch, Toll, Hedgehog, Ras, CCK8 and other secreted entities, involving suites of proteases and enzymes with other functionality – the production of thyroid hormones from transamidated thyroglobulin being especially baroque (Walsh, 2005). Chemokines are examples of complexly derived protein ligands of about 70 aa (Sakmar & Huber, 2016).

Regarded as a pharmacon (bioactive molecule) SgII-70 has as a conserved pharmacophore (active face) **<u>M</u>·<u>KPV/NF</u>·<u>N</u>**. In sheep, with UniProt numbering, the proposed active face is the triply non-contiguous 367**<u>M</u>·**373**<u>KPV</u>**375**<u>F</u>**178**·<u>N</u>**236. In the human this would be 368**<u>M</u>·**374**<u>KPN</u>**376**<u>F</u>**179**·<u>N</u>**237, with N for sheep V. The ovine and human active faces have been deployed here successfully as linear hexapeptides to inhibit prostate cancer cell proliferation in vitro. MKPVFN (EPL601) does not match any ovine (or human) protein and MKPNFN (EPL600) does not match any human (or ovine) protein (NCBI Blast vs non-redundant protein). Pairs of human-related and ovine-related peptides at different sizes performed similarly to each other in mammalian assays, in the activity order 6mers>14mers>8mers. The active face residues comprise two functional groups of three: MKP and V/NFN. Proteins interact with one another primarily via the side chains of surface amino acids. There is enrichment of certain aa at protein interaction sites as compared with their incidence across protein surfaces generally (Sillerud & Larson, 2005). In the enrichment league’s third and fourth places are M and F, respectively, the former the most prevalent of the aliphatics at interaction sites and the latter an aromatic, all of which are over-represented. No enrichment is seen with the other active face residues, K, P, V and N, yet a bimodularity analysis of bioactive ovine peptides emphasized the importance of KP in the forward module and V in the rearward module.

Peptide structure influences ligand binding for sure, but in complex systems in vitro and in vivo also impacts solubility, permeation, half-life, resistance to hydrolysis and other factors unconsidered here, including altered substrate specificity for enzymes such as proprotein convertase and carboxypeptidase, meaning that changes in peptide effects may reflect a factor independent of its biological activity. Limitations of the research further include the consideration that the peptides have not all been tested together in the same assay system. sSgII-70 has been used to make sense of analytical findings and peptide bioactivity, but its existence is a deduction. The attempt to develop peptide mimetics of the active face of a deduced entity might appear quixotic, were it not for the rewards on offer (tissue mass downregulation and reproductive modulation) and the apparent success of the journey from fortuitous 14mer (EPL001) to honed hexamers (including EPL036 and EPL601), for other researchers to explore. The SgII-70 model that has emerged from peptide mimetics is ‘a subset of the six active face residues in appropriate topological conformity will work’. In terms of a ligand-receptor lock-and-key model what is envisaged is a ligand ‘key’ with six ridges, organised in two stereospecific groups of three (MKP & VFN). Two of these keys at the same time access a receptor ‘double lock’ with plenty of give in it. The lock is possibly a granin-style non-canonical receptor, representing a novel protein target for drug development. EPL001 has been deployed in mammalian binding studies to inconclusive effect (Davies et al, 2015). The use of the more reliably active EPL120 and EPL601 can be suggested as the next step in the research, to snare the receptor and enable its deployment to trap the endogenous ligand. Note that receptor binding is proposed to involve both sSgII-70’s amino-terminal MKPVF and a carboxy-terminal N68, acting in concert. For peptide ligands of G-protein coupled receptors it is mostly the C-terminus of the ligand that associates with the transmembrane protein.

In the view articulated here, the 14mers EPL001 and EPL030 are both partial proxies of an endogenous neuroendocrine factor, sSgII-70, and this is why for example supra-hypothalamic administration of either of these anagrammatical tetradecapeptides into the third ventricle of the sheep induces the same pattern of changes in circulating anterior pituitary hormones. (The null hypothesis might speak of shared non-specificity.) The curious situation supervenes of our inferring a hormone’s active face ahead of the hormone’s formal substantiation. Indeed, the potential non-existence of this secreted polypeptide need not hinder the deployment of bioactive peptides; hypothesis-driven research can have useful outcomes even when the hypothesis is wrong. However, the evidence marshalled in this paper does not support the null hypothesis of SgII-unrelatedness. A first expectation, that EPL143, being more faithful to the sSgII-related endogenous entity, is likely to be more active than EPL001, has been upheld (Fig. 5). The second expectation, that concentrating the active face residues as the hexapeptide EPL601 should bolster activity over the 14mer EPL143 has also been upheld. There are 238 atoms present in EPL001 (RPN) and therefore 28,203 interatomic distances (S2). The 15 measurements made here across 21 peptides, although a scintilla of the potential dataset (0.05% in the case of EPL001), have yet delivered meaningful explanations within the concept of SgII- relatedness. The chi-squared probability of EPL001 being dissimilar to the SgII benchmark

EPL143, upholding the null hypothesis, is an artefact of active face residue spacing (see Results, Molecular Modelling). In comparing the sequences of EPL001 and EPL143, Spearman’s rank correlation is 0.88, denoting a very strong positive correlation (S2). The likelihood of an anagram of EPL143, such as EPL001, having the putative active face residues in the same order is 1 in 7,692 (S2). The activity data presented indicate that the SgII-related 6mers EPL600 and EPL601 are useful reductions of the unreliable 14mer EPL001 (inefficacious in the mouse xenograft model, for example, and warranting no further work) and of the active but pharmaceutically impractical 20mer EPL120 (not to mention the pharmaceutically impossible SgII-70). There is a marked improvement in lipophilicity through the series (RPN), using the water/octanol partition coefficient calculated by the Moriguchi method, where Log P should be less than 5 for drug bioavailability: EPL001 = 5.75; EPL120 = 6.35; EPL600 = 0.52; & EPL601 = 1.52; with the reproductive activator EPL036 at 0.60 and the reproductive inactivator EPL630 at 0.21.

## CONCLUSION

Identified by chemical sequencing in the course of an inhibitory hormone discovery project using purified ovine materials, the 14-residue aa sequence designated EPL001, MKPLTGKVKEFNNI, has proved to be deeply encoded within the mammalian genome, but retrievably so it is argued here, in the form of the deduced hormonal factor sSgII-70, using its parent sSgII as the decryption key. Hexapeptides are provided for potential use in reproductive activation (EPL036: IEPVFT, an antagonist of a hormonal OFF signal, delivering reproductive *disinhibition*) and reproductive inactivation (EPL630: KLKMNG, an inhibitory mimetic), and for suppression in cell overgrowth conditions (EPL600 & EPL601: MKPNFN & MKPVFN, inhibitory agonists both).

## AUTHOR CONTRIBUTIONS

JEH conceived and managed the project and wrote the paper, with co-authorial input, notably from RPN and KGD; SM conducted the nematode studies, with KGD; DRH contributed widely to the reporting, analysis and graphical representation of data, including for the supplementary information files; bone marrow cell assays were carried out by BF; mammalian studies in vivo were performed by IJC (sheep) and SDS (mouse); the peptide immobilisation studies were conducted in the laboratory of JAH; bioinformatic analyses were the work of CRM, a ‘red team’ sceptic; molecular modelling was by RPN; all authors approved the work for publication.

## ACKNOWLEDGEMENTS

SM thanks Akanksh Upadhyay for assistance with the nematode studies. Gail Risbridger, in whose laboratory was conducted the prostate cancer cell assay involving ^3^H-thymidine incorporation is thanked for gracious input. SDS thanks Patricia Cooper and Mike Bibby for assistance with the mouse studies. JAH thanks Fanrong Pu for the peptide immobilisation work and Eve Rogers for gene studies. JEH thanks Pat Barker, formerly of The Babraham Institute, Cambridge, UK, and Charles Dickerson, formerly of Harwell Laboratory, South Oxfordshire, UK, for their help in assembling the sequencing and purification record, S1. Similarly, JEH thanks Will Mawby, formerly of the University of Bristol, Bristol, UK, for his help in relation to the Edman sequencing file, S8, and Dave Copsey for myriad contributions to the project.

## SUPPLEMARY INFORMATION (S1-10)

S1, Sequencing & Purification. Figshare: Edman sequencing of biologically active fractions of sheep ovarian follicular fluid and blood plasma. https://doi.org/10.6084/m9.figshare.16432833. This project contains amino acid sequencing data and details how EPL001 and Candidate 7500 were found.

S2, Statistics. Figshare: Statistical analyses applied to data from amino acid sequencing and molecular modelling. https://doi.org/10.6084/m9.figshare.16437891. This project contains the paper’s statistical analyses.

S3, Nematodes. Figshare: Species confirmation of the entomopathogenic nematode, *Steinernema siamkayai,* and pilot study. https://doi.org/10.6084/m9.figshare.16438062. This project contains data confirming the identity of the nematode species used and shows that the results of a pilot study of peptide administration were in line with those of the nematode experiment reported in the paper.

S4, Hypothalamus & Pituitary. Figshare: Histological labelling of the ovine median eminence by an anti-EPL001 antibody and the effects of EPL001 and its scrambled-sequence control EPL030 on pituitary hormone release. https://doi.org/10.6084/m9.figshare.16438161. This project contains raw IHC images showing hypothalamic localisation, together with data on peptide effects on pituitary hormone release in vivo and in vitro.

S5, Murine Studies. Figshare: The toxicity and efficacy of EPL001 and its scrambled-sequence control EPL030 in immunodeficient mice. https://doi.org/10.6084/m9.figshare.16438338. This project presents data showing the lack of toxicity and efficacy of the peptides in immunodeficient mice.

S6, Peptide Immobilisation & Gene Studies. Figshare: The effects of EPL001 on stem cell proliferation, phenotype and gene expression. https://doi.org/10.6084/m9.figshare.16438392. This project presents data showing that EPL001 reduces proliferation and influences gene expression without changing stem cell phenotype.

S7, Interatomic Distances. Figshare: Interatomic distances within peptides related to EPL001. https://doi.org/10.6084/m9.figshare.16438440. This project shows the analysis of 15 interatomic distances between 6 atoms within 21 peptides related to EPL001.

S8, Edman Nonsequentialism. Figshare: Proposed ‘route’ taken by Edman sequencing to provide EPL001 from sSgII-70. https://doi.org/10.6084/m9.figshare.16438452. This project explores the concept of Edman Nonsequentalism, whereby chemical analysis is deemed to have misread the amino acid sequence of the deduced endogenous factor sSgII-70.

S9, Arrive 2.0 Sheep. Figshare: https://doi.org/10.6084/m9.figshare.18480404. This project is a checklist for the ovine study in vivo.

S10, Arrive 2.0 Mouse. Figshare: https://doi.org/10.6084/m9.figshare.18480407. This project is a checklist for the murine studies in vivo.

## INDEPENDENT PEER REVIEW (reprinted with permission)

### Reviewer (Joseph Banoub)

#### Basic reporting

In this manuscript, the authors present an extensive biopurification study aimed to reveal an evolutionarily conserved inhibitory reproductive hormone involved in tissue mass determination. For this reason, they examined the reproduction of the nematode Steinernema siamkayai following fermentation in a system supplemented with different concentrations of exogenous hexapeptides.

The authors used a (rat) bioassay which permitted the physicochemical fractionation (using ovine materials) and Edman degradation, which established the presence of a 14-residue amino acid peptide.

#### Authors ’ re s ponse

No. The presence was established of an ∼70-residue amino acid peptide. This was subject to Edman degradation, providing a 14-residue N-terminal sequence. This was synthesized as the 14-mer synthetic peptide EPL001. **This distinction between the Edman sequence result (14 residues) and the much longer native molecule (∼70 residues) has now been clarified, below.**

Also, they found that the 14-mer synthetic peptide (EPL001), MKPLTGKVKEFNNI, displayed anti-proliferative and reproduction-modulating activity. Moreover, they unexpectedly found that the scrambled-sequence control peptide (EPL030) did likewise.

In addition, they also studied a series of hexapeptides for potential uses for the following: a) in reproductive activation (EPL036: IEPVFT), b) as an antagonist of a hormonal OFF signal, dc) for delivering reproductive disinhibition) and reproductive inactivation (EPL630: KLKMNG, d) as an inhibitory mimetic), and e) for suppression in cell overgrowth conditions (EPL600 & EPL601: MKPNFN & MKPVFN, inhibitory agonists. Furthermore, they established that reproduction increased (x3) after exposure to one synthetic peptide (IEPVFT), while fecundity was reduced (x0.5) after exposure to another (KLKMNG), both effects being dose dependent. They also used hexamers with opposite ends of the synthetic peptide KLKMNGKNIEPVFT (EPL030).

Additionally, they established that EPL001 possessed anti-proliferative effects on human prostate cancer cells and on the rat bone marrow cells. Also, the intra-cerebro-ventricular infusion of EPL001 in sheep showed elevated growth hormone in peripheral blood and reduced prolactin. The highly dissimilar EPL001 and EPL030 nonetheless had similar biological effects in common in mammalian systems, whereas divergently anti-fecundity respectively in the nematode Caenorhabditis elegans.

Peptides up to a 20mer were shown to inhibit the proliferation of human cancer and other mammalian cells in vitro, with reproductive upregulation demonstrated previously in fish and frogs, as well as nematodes.

EPL001was also found to encodes the sheep neuroendocrine prohormone secretogranin II (sSgII). This was deduced on the basis of immunoprecipitation using an anti-EPL001 antibody. And six sSgII residues were responsible for the EPL001bioactivity.

Finally, the authors describe a stereospecific bimodular tri-residue signature involving simultaneous accessibility for binding of the side chains of two specific trios of amino acids, MKP & VFN. An evolutionarily conserved receptor was proposed as having dimeric binding sites, each with ligand-matching bimodular stereocentres.

The bioactivity of the 14mer control peptide EPL030 and its hexapeptide progeny is due to the fortuitous assembly of subsets of the novel hormonal motif, MKPVFN, a default reproductive and tissue building OFF signal.

This manuscript is very well written in Literarily English. It was so well written that this referee found himself wondering what the authors were trying to say. The Introduction part provides an adequate scientific background to pinpoint the aim of this work, the literature references are sufficient.

Although, this referee finds this manuscript long, this is not necessarily a fault, as this length is due to the very comprehensive body of work presented. The article structure, figures, tables and raw data shared are first class.

A quick perusal of the KEY POINTS presented with the manuscript indicate that they are relevant, meaningful and performed to a high technical standard.

The following are some examples that this referee found obscure.

1. In your Abstract, you have written the following:

Background. A biopurification campaign aimed to disclose an evolutionarily conserved inhibitory reproductive hormone involved in tissue mass determination.

Query: What was written makes no sense. I presume the biopurification campaign was started. Also the term campaign is pedantic! As a matter of facts, the whole text is written is Shakespearean style.

#### Authors ’ re s ponse

This has been changed to:

Biopurification has been used to disclose an evolutionarily conserved inhibitory reproductive hormone involved in tissue mass determination.

-Still in your Abstract your Abstract, you have also written the following:

Bioactivity surprises as EPL030 is a scrambled-sequence control of the test peptide, MKPLTGKVKEFNNI (EPL001)

Query: What do you mean by “Bioactivity surprises”?

#### Authors ’ re s ponse

This has been changed to:

Bioactivity is unexpected as EPL030 is a control compound, based on a scrambled sequence of the test peptide MKPLTGKVKEFNNI (EPL001).

-Still in your abstract,

What do you mean when you write that “Both EPL030 and EPL001 are both bioinformatically obscure”?

#### Authors ’ re s ponse

This has been changed to:

EPL030 and EPL001 are both bioinformatically obscure, having no convincing matches to aa sequences in the protein databases.

2. In your Introduction, lines 140-141, you have written the following:

Resistant to conventional bioinformatics analysis (e.g. BLAST searches) and molecular biology, EPL001 is not an incomprehensible singularity: it is an incomprehensible plurality.

Query: I am sure this is written in excellent English grammar, however, I cannot grasp the exact meaning? (Double negative)

#### Authors ’ re s ponse

This has been changed to:

Resistant to conventional bioinformatics analysis (e.g. BLAST searches) and molecular biology, EPL001 is more than an incomprehensible singularity: it is an incomprehensible plurality, having been detected multiple times using Edman degradation (S1).

-Introduction, Lines 166-167, you have written the following:

EPL001 was determined in the context of a polypeptide candidate of m/z ∼7,500 (‘Candidate 166 7500’) in MALDI-TOF mass spectrometry (MS) and a corresponding band in SDS 167 polyacrylamide gel electrophoresis (Hart, 2008), implying a chain of ∼70 aa.

Query: What do want to explain here? EPL001 a polypeptide candidate of m/z ∼7,500 (‘Candidate 166 7500’) in MALDI-TOF mass spectrometry (MS)?

Or are you implying that the protein separated had m/z 7,500? Your test peptide EPL001 has a M.Wt. of 1619 Da

#### Authors ’ re s ponse

This h a s b e en cha ng ed a s fo llo ws, wh ich cla rifie s t h e reviewe r’s m isu n de rstand ing u n de r ‘Ba sic Repo rtin g’:

The 14-residue N-terminal Edman sequence EPL001 was determined in the context of a polypeptide candidate o f m /z ∼7, 5 00 (‘Can dida te 7 5 00 ’) in MAL DI -TOF mass spectrometry (MS) and a corresponding band in SDS 167 polyacrylamide gel electrophoresis (Hart, 2008), implying a chain of ∼70 aa for the native molecule.

-Evinced? Reveal the presence of (a quality or feeling)

#### Authors ’ response

(Change to Line 180.)

The likelihood of a project-unrelated entity having both the sought-for activities is low, especially when evinced

#### Experimental design

The experimental design of the research presented in this manuscript, followed a solid and well thought objectives that were controlled and which permitted and maximized the specific conclusions regarding the following hypothesis statements

-Rat bioassay which permitted the physicochemical fractionation (using ovine materials),

- Edman degradation and state-of-the art mass spectrometry., which established the presence of a 14-residue amino acid peptide.

-The 14-mer synthetic peptide (EPL001), MKPLTGKVKEFNNI, displayed anti-proliferative and reproduction-modulating activity. Moreover, tthe scrambled-sequence control peptide (EPL030) did likewise.

I-The study of a series of hexapeptides for potential uses for the following:

a. in reproductive activation (EPL036: IEPVFT),
b. as an antagonist of a hormonal OFF signal,
c. for delivering reproductive disinhibition) and reproductive inactivation (EPL630: KLKMNG,
d. as an inhibitory mimetic), and
e. for suppression in cell overgrowth conditions (EPL600 & EPL601: MKPNFN & MKPVFN, inhibitory agonists.

- Reproduction increased (x3) after exposure to one synthetic peptide (IEPVFT), while fecundity was reduced (x0.5) after exposure to another (KLKMNG), both effects being dose dependent. They also used hexamers with opposite ends of the synthetic peptide KLKMNGKNIEPVFT (EPL030).

#### Validity of the findings

This manuscript validity can be judged by the results obtained from this research. Indeed, the results obtained are accurate according to the researcher explanation, and prediction. These could be surmised as follows:

The purified ovine materials, the 14-residue aa sequence designated EPL001, MKPLTGKVKEFNNI, has proved to be deeply encoded within the mammalian genome.

The hexapeptides could be used in reproductive activation (EPL036: IEPVFT an antagonist of a hormonal OFF signal, delivering reproductive disinhibition) and reproductive inactivation (EPL630: KLKMNG, an inhibitory mimetic), and for suppression in cell overgrowth conditions (EPL600 & EPL601: MKPNFN & MKPVFN, inhibitory agonists both

#### Additional comments

This manuscript is very well written in Literarily English. It was so well written that this referee found himself wondering what the authors were trying to say. The Introduction part provides an adequate scientific background to pinpoint the aim of this work, the literature references are sufficient.

Although, this referee finds this manuscript long, this is not necessarily a fault, as this length is due to the very comprehensive body of work presented. The article structure, figures, tables and raw data shared are first class.

