## Supplementary Material 2 for "Hexapeptides from a mammalian inhibitory hormone activate and inactivate nematode reproduction"

**Statistics**

A) **Likelihood of a pair of pairs**

In sheep secretogranin II’s second sorting domain there is to be found _367_**ML**K**TG**E**KP**V_375_ (with signal sequence included in residue numbering). Call this string sSgII-9. It is a lightly shuffled version of EPL001. The emphasized residues in sSgII-9 are a match for those from the front half of EPL001, in the form of three doubletons (two being exact matches to EPL001: M**KP**L**TG**KVK), interleaved by singleton matches to the second half of EPL001. The likelihood of a pair of pairs occurring by chance in two 9mer sequences is as follows, where P = probability. P(T) = 1/9. P(TG) = 1/9 x 1/8 = 1/72. TG can be in 8 positions, so 1/72 x 8 = 1/9. P(K) = 2/9. P(KP) = 2/9 x 1/8 = 2/72. KP can be in 8 positions, so 2/72 x 8 = 16/72 = 2/9. P(TG + KP) = 1/9 x 2/9 = 2/81. The probability that TG and KP occur in both sSgII-9 and the EPL001 9mer = (2/81)^2^ = 4/6561 or 1 in 1640.

B) **Lysine placements in Fig. 1 sequence grid**

Residue placement on the sequence grid of the paper’s Fig. 1 is only an issue in regard to EPL001’s three lysines. Where should EPL001’s K2, K7 and K9 be in relation to sSgII-14? There are six ways of positioning three objects, but physical modelling and a couple of assumptions shows that the Fig. 1 arrangement as presented is the only one that ‘works’. EPL001’s KP and TG are present as doubletons in sSgII’s second sorting domain. T & G are adjacent on the grid, so K & P are placed contiguously too. That reduces the six lysine options to two. These are reduced to one by assuming that the stray F & K need to be close enough together probably to be covalently bonded, as they are productively analysed to be in the Discussion. So, sSgII-14’s K11 has to be EPL001’s K9 and not EPL001’s K7. The resulting grid predicts the shape of sSgII-14’s first nine residues (‘sSgII-9’) in free space when modelled in silico (Fig. 1, lower panel).

**C EPL001 versus EPL030: Spearman’s**

The relevant sequences may be compared using Spearman’s rank correlation (r_s_). If the EPL001 residues MKPLTGKVKEFNNI are numbered 1-14, they appear in EPL030,

KLKMNGKNIEPVFT, in the order 2, 4, 9, 1, 12, 6, 7, 13, 14, 10, 3, 8, 11 and 5, selecting the most adjacent Ks and Ns. The difference, d, is 2, 2, 6, -3, 7, 0, 0, 5, 5, 0, -8, -4, -2, -9 and thus d^2^ is 4, 4, 36, 9, 49, 0, 0, 25, 25, 0, 64, 16, 4, 81, with the sum (∑) of d^2^ equalling 317. Thus: r_s_ = 1 – (6∑d^2^)/n(n^2^-1) = 0.30, denoting a weak positive correlation. Randomisation of the test peptide EPL001 to provide EPL030 as a scrambled-sequence control was therefore achieved, albeit sub-optimally, as described in the paper’s Discussion.

**Interatomic distances within EPL001**

There are 238 atoms present in the 14mer peptide EPL001 (RPN). Between atom 238 and the rest there are 237 distances. Between atom 237 and the rest, excluding atom 238, there are 236 distances. Thus the distances available are 237+236+235...+3+2+1, i.e. the sum of factorial 237. Calculate according to Gauss: +!n = n(n+1)/2 = 237(237+1)/2 = 28,203.

As a check, the number series may be written out in arithmetic order in one row and in reverse order in the row below it:

1	2	3	… 235 236 237

237 236 235 …	3	2	1

Reading vertically each column of two numbers sums to 238.

There are 237 columns in all, so

238 x 237 = 56,406

But we have two rows and must divide by two to get the sum of the numbers from 1 to 237:

56,406/2 = 28,203

The interatomic distance measurements made in the present study on EPL001 were 15 in all, representing 0.05% of the potential dataset.

**Tri-residue analysis as applied to EPL001**

Using molecular modelling, aa residues in 20 peptides were evaluated visually in groups of three and viewed clockwise and counter clockwise, to provide a spatial assessment of tri-residue aa side chain availability for receptor binding interactions (see Molecular Modelling, Materials and Methods).

EPL001 is a 14mer peptide of aa sequence MKPLTGKVKEFNNI. The number of ways of selecting 3 items from 14 is expressed by the equation n!/r!(n-r)!, where n is 14 and r is 3. n! = 14! = 87,178,291,200. r! = 3 x 2 x 1 = 6. (n-r)! = (14-3)! = 11! = 39,916,800. So: 87,178,291,200/6 x 39,916,800 = 87,178,291,200/239,500,800 = 364. The 364 tri-residue groups can be viewed such that they are clockwise in the modelling image or counter clockwise.

Of the 364 tri-residue combinations in EPL001, those relating to 6 residues in EPL001’s 14 aa were the focus of the analysis. These residues are the aa postulated to comprise an SgII-related hormonal active face: M, K2, P, V, F & N12. The number of ways of selecting 3 items from 6 is n!/r!(n-r)!, where n is 6 and r is 3. So: 6 x 5 x 4 x 3 x 2 x 1/3 x 2 x 1 (6-3)! = 6 x 5 x 4 x 3 x 2 x 1/3 x 2 x 1 (3 x 2 x 1) = 720/36 = 20. Each of these 20 combinations of three can be viewed via modelling to provide clockwise and counter clockwise images. Of EPL001’s total tally of tri-residue combinations of 364 the 20 active face permutations represent 5.5%.

**EPL143 versus EPL001: Spearman’s**

The relevant sequences may be compared using Spearman’s rank correlation (r_s_). If the rank order of sSgII-14 residues (MLKTGEKPV∙FK∙NNI) in the contiguous sequence EPL143 (MLKTGEKPVFKNNI) is 1-14 then by comparison those in EPL001 (MKPLTGKVKEFNNI) are 1, 4, 2, 5, 6, 10, 7 ,3, 8, 11, 9, 12, 13, 14, selecting the most adjacent Ks and Ns. The difference, d, is 0, -2, 1, -1, -1, -4, 0, 5, 1, -1, 2, 0, 0, 0 and thus d^2^ is 0, 4, 1, 1, 1, 16 (non-matching E), 0, 25 (non-matching P), 1, 1, 4, 0, 0, 0, with the sum (∑) of d^2^ equalling 54. Thus: r_s_ = 1 – (6∑d^2^)/n(n^2^-1) = 0.88, denoting a very strong positive correlation.

**Anagram of EPL143/sSgII-14 having postulated active face residues in the same order**

***SOURCE SEQUENCE:***

MLKTGEKPVFKNNI ( = EPL143, being the synthesized contiguous form of sSgII-14 from the paper’s Fig. 1)

***ANAGRAM:***

MKPLTGKVKEFNNI ( = EPL001, a synthesized 14mer peptide)

These two sequences can be represented with emphasis on the postulated active face residues for receptor binding:

**M**LKTGE**KPVF**K**N**NI

**MKP**LTGK**V**KE**FN**NI

The active face residues are in the same order in this pair of anagrams. Starting with EPL143, what is the chance of obtaining an anagram of it, such as EPL001, having the postulated active face residues in the same order, though not necessarily in the same position?

The number of anagrams of a 14mer such as EPL143 is 14! = 87.18bn. How many of these have the active face residues in the right order, i.e. MKPVFN?

The starting case is this:

MKPVFNxxxxxxxx

Imagine a bag of 14 balls representing the residues in EPL143. The likelihood of picking each one of these balls in turn in the right order from the bag to get MKPVFNxxxxxxxx

is as follows:

1/14 x 3/13 (because there are three Ks in the EPL143/sSgII-14 source sequence) x 1/12 x 1/11 x 1/10 x 2/9 (because there are two Ns)

Each residue can be in one of 8 positions, so:

8 x 6 = 48

The calculation is therefore:

1 x 3 x 1 x 1 x 1 x 2 x 48/14 x 13 x 12 x 11 x 10 x 9 = 288/2,162,160 = 1.3 x 10-4 = 0.00013

So P = 0.00013, i.e. the probability of pulling an in-order active face 14mer like EPL001 out of the bag of EPL143 is that number. To put it another way there is a 13 in 100,000 chance of pulling an MKPVFN-containing peptide with any gapping from the EPL143/sSgII-14 bag, which is 1 in 7,692.

**Statistical support for Supplementary Information 1, Sequencing & Purification (S1)**

The doubleton PV is present in (i) the First Sighting (SEQ ID NO: 1) as Mxxxx**PV**, (ii) the second sorting domain of sSgII as Mxxxxx**PV** and (iii) the homologue of the sSgII second sorting domain in the fruit fly protein Q9W2X8 as Mxxxxxxxxxxx**PV**, with 11 intervening residues. The theoretical probability of obtaining the First Sighting, MxxxxPVxxxxxxx, by chance from the universe of potential 14 aa sequences is 1/20 x 20/20 x 20/20 x 20/20 x 20/20 x 1/20 x 1/20 x 20/20 x 20/20 x 20/20 x 20/20 x 20/20 x 20/20 x 20/20 = 1/20 x 1/20 x 1/20. But there are 12 possible positions for PV, given an initial M. The theoretical probability thus becomes 1/20 x 1/20 x 1/20 x 12 = 0.0015 or 1 in 666. The actual situation can now be considered. In the SwissProt database (accessed via UniProt) there are 547,085 proteins, with a total of ~195m aa. The incidence of each of the 12 different M-PV forms, from 3mer up to 14mer, is within the range 14,000-17,000 (author CRM). The number of overlapping mers in the database ranges from ~194m 3mers down to ~188m 14mers. Dividing each of these numbers by the number of M-PVs in each category delivers a probability in each case of 0.000074-0.000087, i.e. 1 in 11,400-13,500. Summing these yields a total probability of 0.000993. The chance of selecting a single sequence comprising M-PV, where the dash represents 0-11 residues, is therefore 1 in 1007. There is thus only a small chance of the First Sighting sequence containing M-PV at random.

**Statistical support for Supplementary Information 8, Edman Nonsequentialism (S8)**

To get from sSgII-14 to EPL001 requires to be moved only five sSgII-14 residues, four charged and one in a class of its own: K3**^+^**, K7**^+^**, K11**^+^**, E6**^-^** and P8 (unique in being a proteinogenic *secondary* amino acid). The probability of choosing one uncharged residue of any kind from sSgII-14 and the four charged residues is 10/14 x 4/13 x 3/12 x 2/11 x 1/10 x (5!/(4! x 1!)) = 5/1001 or about 1 in 200. The probability of choosing proline specifically as the uncharged residue, together with the four charged residues is 1/14 x 4/13 x 3/12 x 2/11 x 1/10 x (5!/(4! x 1!)) = 1/2002 or about 1 in 2000. These calculations support the view that something puzzling went on with the Edman chemical sequencing, relating in part to charged residues.

There are two mixed signals in the Beale 4 Edman reading, xxPxxxx**V/L**xx**F/K**Nxx (SEQ ID NO: 4 in S1 Table 1). The Beale 4’s V & F are in register with the V & F in EPL001 and each of these Beale 4 residues is paired with co-available residues, L & K, having free α amines according to S8 Fig. 1 scheme by which EPL001 is derived from sSgII-14. The probability of getting both V/L & F/K in the same EPL001-related sequence by chance *in any position* is 1/14 x 1/13 x 1/13 x 3/12 (given that there are 3 Ks) x 182 positions = 546/28392 or 1 in 52. (The 182 positions are calculated by reference to number of spaces between the two pairs versus number of possible positions: 0 = 13, 1 = 12, 2 = 11 etc, to 12 = 1, total 91; x2 for V/L preceding F/K and vice versa = 182.) The probability of getting V/L & F/K in the same EPL001-related sequence by chance *specifically in positions 8 and 11* is 1 in 167.
